# Integrin α11 is enriched in quiescence and promotes cell-cycle re-entry through destabilisation of the CDK inhibitor p27

**DOI:** 10.64898/2026.08.29.748043

**Authors:** Ekjot Kaur, Jordan A. Holt, Rona Wilson, Van Kelly, Elena Gomez-Marin, Bojan Zunar, Alasdair Daniels, Rozita Adib, Philipp Thomas, Boris Lenhard, Tony Ly, Alexis R. Barr

## Abstract

Proteins that distinguish quiescent cells from other non-proliferative states and actively regulate their return to proliferation remain poorly understood. Here, we combined quantitative proteomics with functional image-based screening to identify regulators of the quiescence-to-proliferation transition. Amongst the functional quiescence signature proteins we identified, we focussed on integrin α11 (*ITGA11*) which is induced across multiple models of reversible quiescence in distinct cell types and that has low expression in proliferating and senescent cells. Although *ITGA11* is dispensable for proliferation of asynchronously cycling cells, it is required for efficient cell-cycle re-entry from quiescence. Mechanistically, *ITGA11* promotes YAP accumulation and nuclear localization, thereby sustaining *SKP2* expression and p27 degradation during cell cycle re-entry. Depletion of p27, or pharmacological activation of YAP signalling rescues the cell-cycle re-entry defect caused by *ITGA11* depletion. Together, these findings identify *ITGA11* as a functional quiescence signature protein that couples extracellular matrix sensing to YAP-dependent regulation of the Skp2-p27 axis, revealing a mechanism that controls the transition from quiescence to proliferation.

## Introduction

Quiescence (G0) is a reversible, non-proliferative state that enables cells to preserve long-term viability, maintain tissue integrity, and rapidly re-enter the cell cycle in response to mitogenic cues. Quiescence arises in a wide range of physiological contexts, including adult stem cell maintenance, tissue homeostasis, wound repair, DNA damage^1, 2^ and adaptation to fluctuating nutrient and growth factor availability^3^. Experimentally, quiescence can be induced by diverse stimuli such as growth factor deprivation, nutrient limitation, contact inhibition, extracellular matrix signalling, or pharmacological inhibition of Cyclin-CDK activity^3^. Although these stimuli engage distinct upstream pathways, they converge on a common cellular programme characterized by withdrawal from the cell cycle while preserving the capacity for future proliferation.

Unlike senescence, which represents a terminal, and frequently inflammatory, growth arrest, quiescence is defined by low metabolic activity, suppression of cell-cycle gene transcription, and the capacity for rapid reactivation^4, 5^. Entry into and exit from quiescence are tightly regulated by Cyclin-CDK complexes and their inhibitors, particularly p21(Cip1/Waf1) and p27(Kip1), which integrate environmental signals to modulate G1-S progression^6, 7^. Transcription factors including the DREAM complex, FOXO family members, Hes1, and p53-associated regulators (including p21) suppress cell-cycle progression while preserving cellular fitness and the capacity for rapid reactivation^8^. Increasing evidence indicates that quiescence is not a uniform state but comprises distinct programs defined by specific combinations of transcription factors and niche-derived signals, reflecting substantial heterogeneity across tissues and experimental models. Similarly, senescence is not a single homogeneous state but represents a spectrum of stable cell-cycle arrested phenotypes that vary depending on the nature of the inducing stress, cellular context, and tissue of origin. Senescence is triggered by persistent stress, including DNA damage, telomere dysfunction, excessive cell size^9–12^, or oncogenic signalling^13^. Senescent cells exhibit chronic activation of the DNA damage response^14, 15^, p16^INK4a^–dependent RB enforcement^16^, extensive chromatin remodelling including ATRX-associated heterochromatin changes^17^, senescence-associated distension of satellites (SADS) formation^18^ and lamina-associated domains (LADs) detachment^19^, and development of a pro-inflammatory senescence-associated secretory phenotype (SASP)^20^. Despite extensive characterization of senescence maintenance pathways, the mechanisms that allow quiescent cells to retain proliferative competence and the lack of definitive markers distinguishing quiescence from senescence remain major challenges for the field.

The extracellular matrix (ECM) and its cognate receptors, the integrins, play central roles in coordinating cell-cycle entry with mechanical and biochemical features of the microenvironment. Integrins transduce ECM stiffness, composition, and topology into intracellular signalling through focal adhesion kinase (FAK), Rho GTPases, and downstream transcriptional regulators^21, 22^. Classical studies have shown that integrin engagement is required for G1 progression, in part through regulation of Cyclin D1 expression, p27 levels, and cytoskeletal tension^23–25^. However, the specific integrin subunits that govern quiescence maintenance and exit, and the molecular pathways they engage, remain poorly defined.

Here, we sought to determine the molecular features that differentiate quiescence from senescence and to understand how quiescent cells retain their proliferative potential. Using quantitative proteomics followed by phenotypic screens for quiescence-proliferation transitions, we identify proteins that are enriched or depleted in quiescent cells, as compared to senescent and G1 cells, and define those that play a role in quiescence-proliferation transitions. We characterise integrin α11 (encoded by *ITGA11*) as a key integrin involved in promoting cell cycle entry from quiescence and show how integrin α11 influences the mechano-responsive transcription factor, YAP, to regulate p27 stability and cell cycle entry.

Together, this work advances our understanding of reversible cell-cycle arrest and identifies mechanisms that enable quiescent cells to preserve proliferative potential, with implications for tissue homeostasis, regeneration, ageing, and disease.

## Results

### Enrichment of proteins governing quiescence

To delineate the molecular pathways distinguishing reversible quiescence from irreversible senescence, we performed quantitative proteomics on G1, quiescent and senescent hTert-RPE1 mRuby-PCNA p21-GFP cells. We used FACS to isolate three states of hTert-RPE1 cells expressing endogenously tagged mRuby-PCNA and p21-GFP^1^: p21Low PCNALow G1 cells, p21High PCNALow spontaneously quiescent cells (where cells enter quiescence in response to intrinsic DNA damage^1, 2^), and p21High senescent cells (exogenous (etoposide) DNA damage-induced) (Supplementary Fig. 1A-C). Differential protein expression analysis and unsupervised clustering of the total proteome resolved 148 proteins enriched or depleted in quiescent and/or senescent populations (Fig. 1A; Supplementary Tables 1-2). Gene ontology analysis revealed that these 148 proteins were enriched into 12 molecular function categories, encompassing regulators of extracellular matrix (ECM) organization, cyclin dependent kinase (CDK) activity, and metabolic processes (Supplementary Fig. 1D). This enrichment analysis suggested a coordinated rewiring of multiple cellular systems during cell-cycle exit.

**Figure 1:**
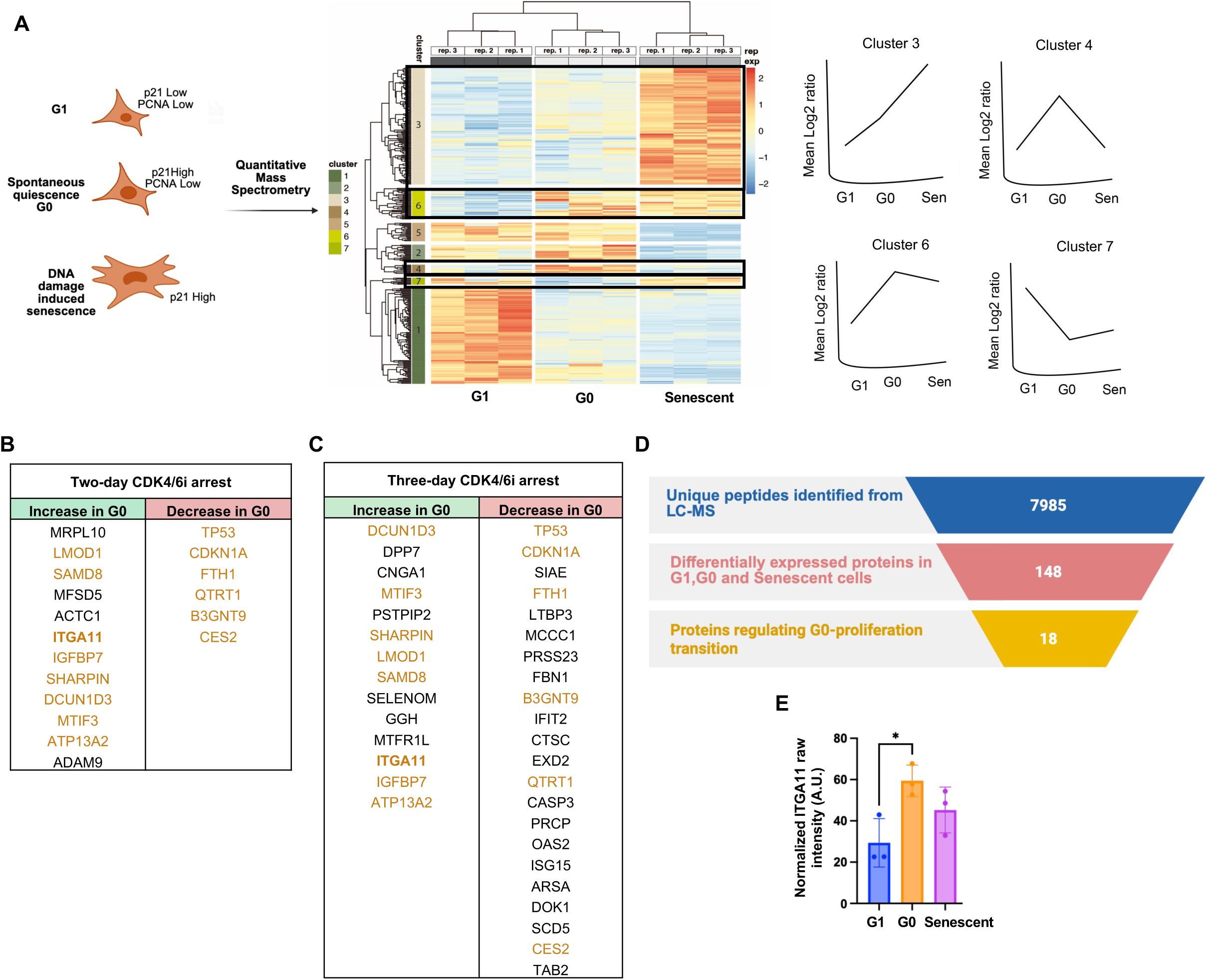
Enrichment of proteins governing quiescence. A) Schematic representation of fluorescence-activated cell sorting (FACS) of cells, based on endogenous p21-GFP and mRuby-PCNA expression, for mass spectrometry. Unsupervised clustering of quantitative proteomic data from p21Low PCNALow (G1), p21High PCNALow (G0) and p21High (senescent) hTert-RPE1 mRuby-PCNA p21-GFP cells. Protein intensities are color-coded from blue (decreased) to red (increased), as indicated in the key. Cluster 3 comprises proteins enriched in senescent cells, whereas clusters 4 and 6 contain proteins enriched in quiescent cells. Cluster 7 contains proteins depleted in quiescent cells relative to G1 and senescent cells. Three biological replicates were analysed per condition.B) Summary of candidate genes identified from the two-day CDK4/6 inhibitor (CDK4/6i) arrest and release screen. Gene depletion that increased the G0 fraction (decreased EdU incorporation) is shown in green, whereas depletion that decreased the G0 fraction (increased EdU incorporation) is shown in red. Candidate genes identified in both screening conditions are highlighted in orange. C) Summary of candidate genes identified from the three-day CDK4/6i arrest and release screen. Gene depletion that increased the G0 fraction (decreased EdU incorporation) is shown in green, whereas depletion that decreased the G0 fraction (increased EdU incorporation) is shown in red. Candidate genes identified in both screening conditions are highlighted in orange. D) Quantitative LC-MS analysis identified 7,985 proteins across all samples. Differential expression analysis identified 148 proteins significantly altered in quiescent and/or senescent cells relative to G1 cells. Functional prioritisation and secondary screening identified 18 candidate regulators of the G0-to-proliferation transition. E) Normalized ITGA11 protein abundance measured by quantitative proteomics (cluster 6) in p21Low PCNALow (G1), p21High PCNALow (G0), and p21High senescent hTert-RPE1 mRuby-PCNA p21-GFP cells. Data are presented as mean ± SD from three biological replicates. Statistical significance was determined by one-way ANOVA with multiple-comparison correction. *P < 0.05.

To interrogate whether these proteins influence quiescence entry or exit, we conducted an image-based phenotypic siRNA screen in asynchronously proliferating hTert-RPE1 cells and in cells released from quiescence back into proliferation. We used CDK4/6 inhibition (CDK4/6i, Palbociclib) to arrest cells in quiescence^26^ for either two or three days (Supplementary Fig. 1E-F). Two-and three-days CDK4/6i arrest and release experiments were selected to model fully reversible quiescence (two-day CDK4/6i) and becoming irreversible (three-day CDK4/6i), since beyond two days arrest in CDK4/6i, hTert-RPE1 cells start to enter irreversible senescence^10, 27^. This allowed us to assess if gene depletion altered the ability of cells to enter an irreversible senescent arrest. Quantitative assessment of EdU incorporation demonstrated high reproducibility across technical repeats (Supplementary Fig. 1G-I). For each condition, Z-scores were calculated for the fraction of EdU-positive cells, representing cells that re-entered the cell cycle following release from CDK4/6i-induced quiescence or under asynchronous growth conditions (Supplementary Tables 3-5).

In asynchronously cycling cells, depletion of 14 genes decreased the EdU-positive fraction (Supplementary Table 6). These genes may impact proliferation independently of quiescence and were therefore not considered further here. After removing these 14 genes from consideration, 12 genes impaired cell cycle re-entry from a two-day CDK4/6i arrest (Fig. 1B) and 14 genes impaired cell cycle re-entry from a three-day CDK4/6i arrest (Fig. 1C), suggesting that these genes normally facilitate cell cycle re-entry from quiescence. Conversely, depletion of 6 genes (release from two-day CDK4/6i (Fig. 1B)) and 22 genes (release from three-day CDK4/6i (Fig. 1C)) increased the fraction of cells returning to proliferation after release from quiescence, consistent with these genes acting to normally promote quiescence or senescence, or acting to inhibit cell cycle entry.

Integration of the proteomic signatures with the functional data from the siRNA screen yielded 18 proteins with potential roles in controlling re-entry into the cell cycle from quiescence (Fig. 1D). Among these, integrin α11 (or integrin subunit alpha 11, *ITGA11*), an extracellular matrix-associated integrin subunit, emerged as a robust and previously uncharacterized candidate. Integrin α11 is a collagen-binding integrin highly expressed in fibroblasts and stromal cells, where it contributes to ECM remodelling, tissue repair, and tumour-stroma interactions^28, 29^. Although *ITGA11* has been implicated in fibrosis and cancer-associated fibroblast activation, its potential role in regulating quiescence and quiescence-proliferation transitions have not been explored. Our proteomic analyses revealed that *ITGA11* is selectively enriched in spontaneous quiescence, as compared to G1 and senescent cells (Fig. 1E). Additionally, siRNA-mediated depletion of *ITGA11* by siRNA did not show any effect on asynchronously cycling cells in the screen but caused a decrease in cell cycle re-entry in cells released from a two- or three-day CDK4/6i G0 arrest (Supplementary Fig. 1E-F), raising the possibility that *ITGA11* functions as a previously unrecognized adhesion-dependent regulator of quiescence exit. Notably, *ITGA11* knockout mice exhibit reduced body size, a phenotype consistent with impaired proliferative capacity, although the underlying cellular mechanisms remain undefined^30, 31^. As a cell-surface type I collagen receptor, *ITGA11* is also an attractive candidate for therapeutic targeting, offering the potential to modulate quiescence-associated signalling or selectively deliver therapeutic agents to quiescent cell populations. Together, these observations prompted us to investigate the molecular role of *ITGA11* in regulating cell-cycle re-entry.

### Integrin α11 is upregulated in different types of quiescence

To determine whether the enrichment of *ITGA11* identified in our proteomic analysis of spontaneously quiescent cells reflected a broader quiescence-associated expression change, we assessed integrin α11 protein expression across different models of quiescence. Integrin α11 protein levels were quantified by immunoblotting in spontaneously arising p21high quiescent cells (FACS purified the same way as used for the initial proteomic analysis; Fig. 1A), CDK4/6i-induced quiescent cells and contact-inhibited quiescent cells and compared with asynchronously cycling and etoposide-induced senescent hTert-RPE1 cells. CDK4/6i-induced, contact-inhibited quiescent and etoposide-induced senescent hTert-RPE1 cells exhibited decreased percentage of EdU positive cells compared to the asynchronously growing cells, confirming the establishment of the respective cell states (Supplementary Fig.2A). Across all three quiescence models, integrin α11 protein was consistently elevated relative to proliferating and senescent cells, with protein levels significantly exceeding those observed in senescent populations (Fig. 2A-B). The specificity of the integrin α11 antibody was validated by siRNA-mediated depletion of *ITGA11* in multiple quiescence models, which resulted in a marked reduction in integrin α11 protein levels (Supplementary Fig. 2B-C). These findings indicate that *ITGA11* upregulation is associated with reversible cell-cycle exit and is not simply correlated with p21 expression.

**Figure 2:**
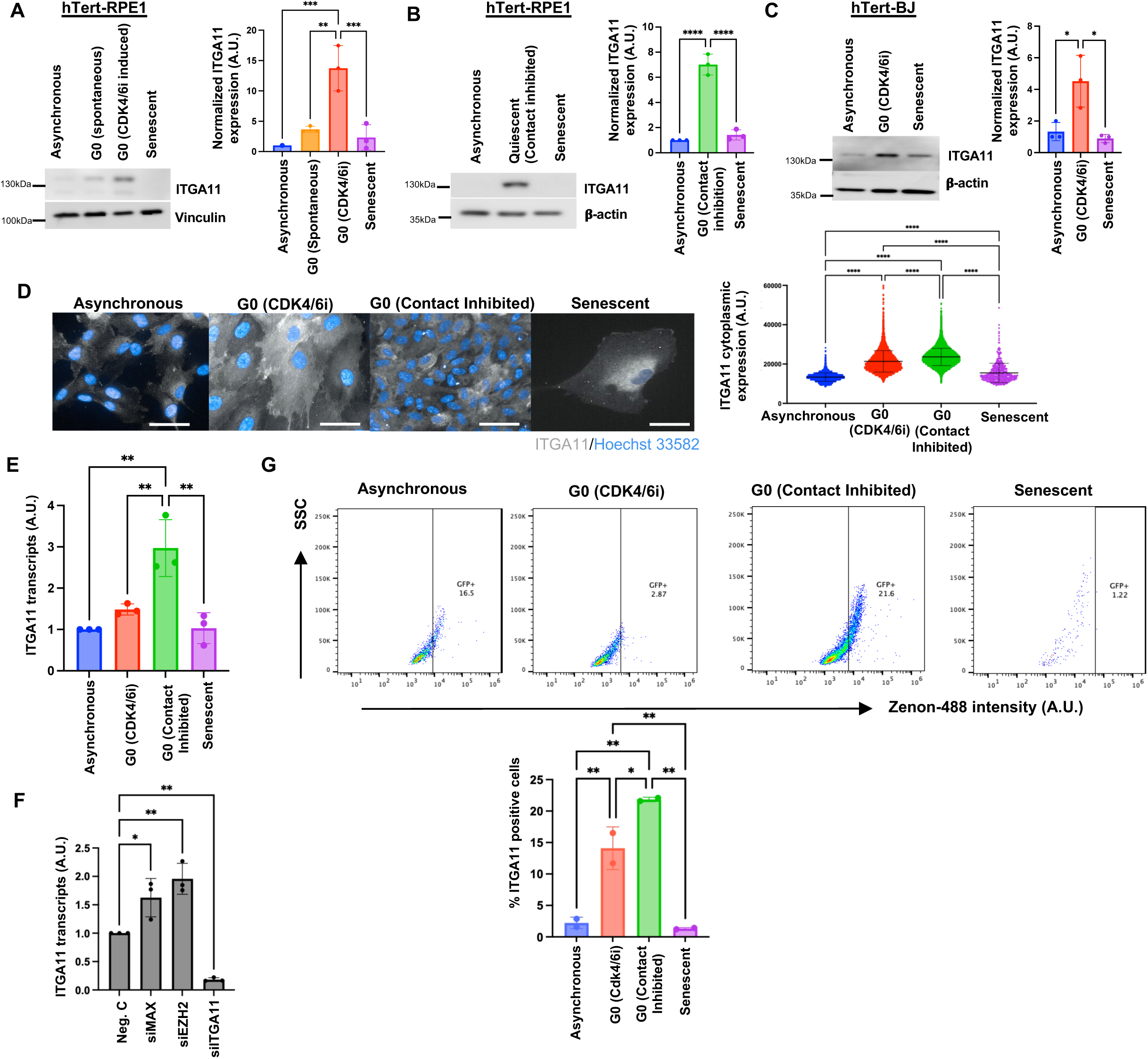
Integrin α11 is upregulated in different types of quiescence. A) (Left) Western blot analysis for integrin a11 in asynchronous, p21High PCNALow spontaneous quiescent (G0), 1µM CDK4/6i-induced quiescence and 50µM etoposide induced senescent hTert-RPE1 mRuby-PCNA p21-GFP cells. Vinculin is used as loading control. (Right) Quantification of the integrin α11 intensity normalized to vinculin. Data are presented as mean ± SD from three biological replicates. Statistical significance was determined by one-way ANOVA with multiple-comparison correction. ***P < 0.001. B) (Left) Western blot analysis of integrin α11 expression in contact inhibition-induced quiescent, asynchronous, and 50μM etoposide-induced senescent hTert-RPE1 cells. β-actin is used as loading control. (Right) Quantification of the integrin α11 intensity normalized to β-actin. Data are presented as mean ± SD from three biological replicates. Statistical significance was determined by one-way ANOVA with multiple-comparison correction. ****P < 0.0001. C) (Left) Western blot analysis of integrin α11 expression in asynchronous hTert-BJ fibroblasts, following induction of quiescence with 1μM CDK4/6 inhibitor and senescence with 50μM etoposide. β-actin is used as loading control. (Right) Quantification of the integrin α11 intensity normalized to β-actin. Data are presented as mean ± SD from three biological replicates. Statistical significance was determined by one-way ANOVA with multiple-comparison correction. *P < 0.05. D) (Left) Representative immunofluorescence images of integrin á11 localization in asynchronous, quiescent, and senescent hTert-RPE1 cells. (Right) Quantification of cytoplasmic integrin á11 fluorescence intensity measured within the cytoplasm segment based on vinculin channel intensity Representative of n=2 biological replicates. Statistical significance was determined by one-way ANOVA with multiple-comparison correction. ****P < 0.0001. Scale bar is 100μm. E) *ITGA11* mRNA expression in asynchronous, quiescent, and senescent cell populations measured by SYBR Green qPCR. *GAPDH* was used as the reference gene. Data are shown as mean ± SD (n = 3 biological replicates). Statistical significance was determined by one-way ANOVA with multiple-comparison correction. **P < 0.01. F) *ITGA11* mRNA expression following siRNA-mediated depletion of *MAX* or *EZH2* for 48hrs. *ITGA11* siRNA was included as a positive control. Statistical significance was determined by one-way ANOVA with multiple-comparison correction. *P < 0.05, ** P < 0.01. G) (Above) Flow cytometric analysis of cell-surface integrin á11 expression using a Zenon™ 488-labelled antibody in cells induced into quiescence by 1µM CDK4/6i or contact inhibition, and cells induced into senescence by treatment with 50ìM etoposide. (Below) Quantification of percentage of Zenon-488 anti-integrin α11 positive cells. Data are shown as mean ± SD (n = 2 biological replicates). Statistical significance was determined by one-way ANOVA with multiple-comparison correction. *P < 0.05, **P < 0.01.

To determine whether this regulation was conserved across non-epithelial cell types, we examined integrin α11 expression in hTert-BJ fibroblasts. CDK4/6i-induced quiescent and etoposide-induced senescent hTert-BJ fibroblasts exhibited a reduced fraction of EdU-positive cells compared with asynchronously proliferating cells, validating the quiescent and senescent phenotypes (Supplementary Fig. 2D). Similar to hTert-RPE1 cells, integrin α11 expression was highest in quiescent hTert-BJ fibroblasts and was markedly lower in proliferating and senescent populations (Fig. 2C), suggesting that integrin α11 upregulation is a conserved feature of cellular quiescence.

We next examined the subcellular distribution of integrin α11 across cell states. Immunostaining revealed enhanced cytoplasmic accumulation of integrin α11 in quiescent cells, with the strongest enrichment observed in CDK4/6i-treated and contact-inhibited populations, consistent with immunoblotting (Fig. 2D). The specificity of the integrin α11 antibody for immunostaining was confirmed using siRNA against *ITGA11* (Supplementary Fig. 2E).

To investigate whether *ITGA11* upregulation occurred at the transcriptional level, *ITGA11* mRNA expression was quantified in asynchronous, quiescent, and senescent hTert-RPE1 cells. Increased *ITGA11* transcript levels were observed in CDK4/6i quiescent and contact-inhibited cells (Fig. 2E). This suggests that transcriptional regulation contributes to *ITGA11* expression in quiescence.

To identify potential regulators of *ITGA11* expression, we interrogated the ENCODE database^32^ and identified 16 candidate transcription factors with predicted binding sites within the *ITGA11* promoter. We performed Super-Low Input Carrier-CAGE (SLIC-CAGE) on cells isolated from distinct cell-cycle states, including G1 and G0 (contact inhibition), to identify transcription factors with cell-state-dependent expression. This analysis revealed differential expression of *MAX* and *EZH2* (Supplementary Fig. 2F). *MAX* is a basic helix-loop-helix/leucine zipper (bHLH/LZ) protein that forms heterodimers with multiple DNA-binding partners, including *c-MYC* and *MAD* family proteins, to regulate gene expression^33^. *EZH2* is a catalytic component of Polycomb repressive complex 2 (PRC2) that methylates histone 3 lysine 27 (H3K27me3), promoting epigenetic silencing of target genes^34^. Compared with G1 cells, G0 cells exhibited increased *MAX* transcript levels and reduced *EZH2* expression (Supplementary Fig. 2F). Analysis of ENCODE ChIP-sequencing datasets showed that *EZH2* binds the *ITGA11* promoter within 5kb upstream of the transcription start site, whereas *MAX* occupies regions downstream of the promoter, as well as sites within the *ITGA11* gene body (Supplementary Fig. 2G). To determine whether these factors regulate *ITGA11* expression, *MAX* and *EZH2* were individually depleted by siRNA in asynchronously proliferating hTert-RPE1 cells (Supplementary Fig. 2H), and *ITGA11* transcript levels were quantified by qPCR. Depletion of either *MAX* or *EZH2* significantly increased *ITGA11* expression in asynchronous cells (Fig. 2F), indicating that both factors normally repress *ITGA11* transcription in actively cycling cells. The increase in *ITGA11* expression following *EZH2* depletion is consistent with its established role as a transcriptional repressor through PRC2-mediated H3K27 trimethylation. Similarly, the induction of *ITGA11* upon *MAX* depletion suggests that *MAX* contributes to maintaining repression of *ITGA11* in proliferating cells, either directly through *MAX-MAD* or other repressor complexes or indirectly through regulation of transcriptional programs associated with cell proliferation.

As integrin α11 is a cell-surface receptor, we next examined whether surface integrin α11 expression could distinguish quiescent from senescent cells, addressing the need for markers that distinguish quiescent from senescent arrested states. Flow cytometric analysis was performed on asynchronous, quiescent (CDK4/6i-induced and contact-inhibited), and senescent hTert-RPE1 cells using a Zenon 488-conjugated integrin α11 antibody. Surface integrin α11-positive cells were markedly enriched in quiescent populations, with 15% and 21.6% positive cells observed in CDK4/6i-induced and contact-inhibited cells, respectively, compared with 2.87% in asynchronous cells and 1.22% in senescent cells (Fig. 2G and Supplementary Fig. 2I). The specificity of the cell surface integrin α11 expression was validated by performing FACS analysis in the Neg.C and siITGA11 transfected CDK4/6i-induced quiescent hTert-RPE1 cells (Supplementary Fig. 2J).

Collectively, these results demonstrate that integrin α11 is selectively upregulated across multiple models of quiescence and in distinct cell types. Furthermore, integrin α11 exhibits increased cell-surface expression in quiescent cells, supporting its utility as a marker of reversible, quiescent cell-cycle arrest and suggesting a role in quiescence-associated adhesion and signalling pathways.

### *ITGA11* promotes cell-cycle re-entry following quiescence

To define and validate the functional contribution of *ITGA11* to the quiescence-proliferation transition, we depleted *ITGA11* using either different siRNA pools (siRNA Tools) to those used in our original screen (Supplementary Fig. 2E) or doxycycline-inducible shRNA (Supplementary Fig. 3A). Validating our original screen, these independent approaches to deplete *ITGA11* significantly decreased the fraction of cells returning to proliferation following release from CDK4/6i-induced quiescence (Fig. 3A, Supplementary Fig. 3B) whereas asynchronously cycling cells were largely unaffected (Fig. 3B, Supplementary Fig. 3C). Additionally, *ITGA11* depletion produced an increase in cell clustering amongst cells 24hrs post CDK4/6i-release, a phenotype absent in negative-control siRNA conditions (Supplementary Fig. 4A)

**Figure 3:**
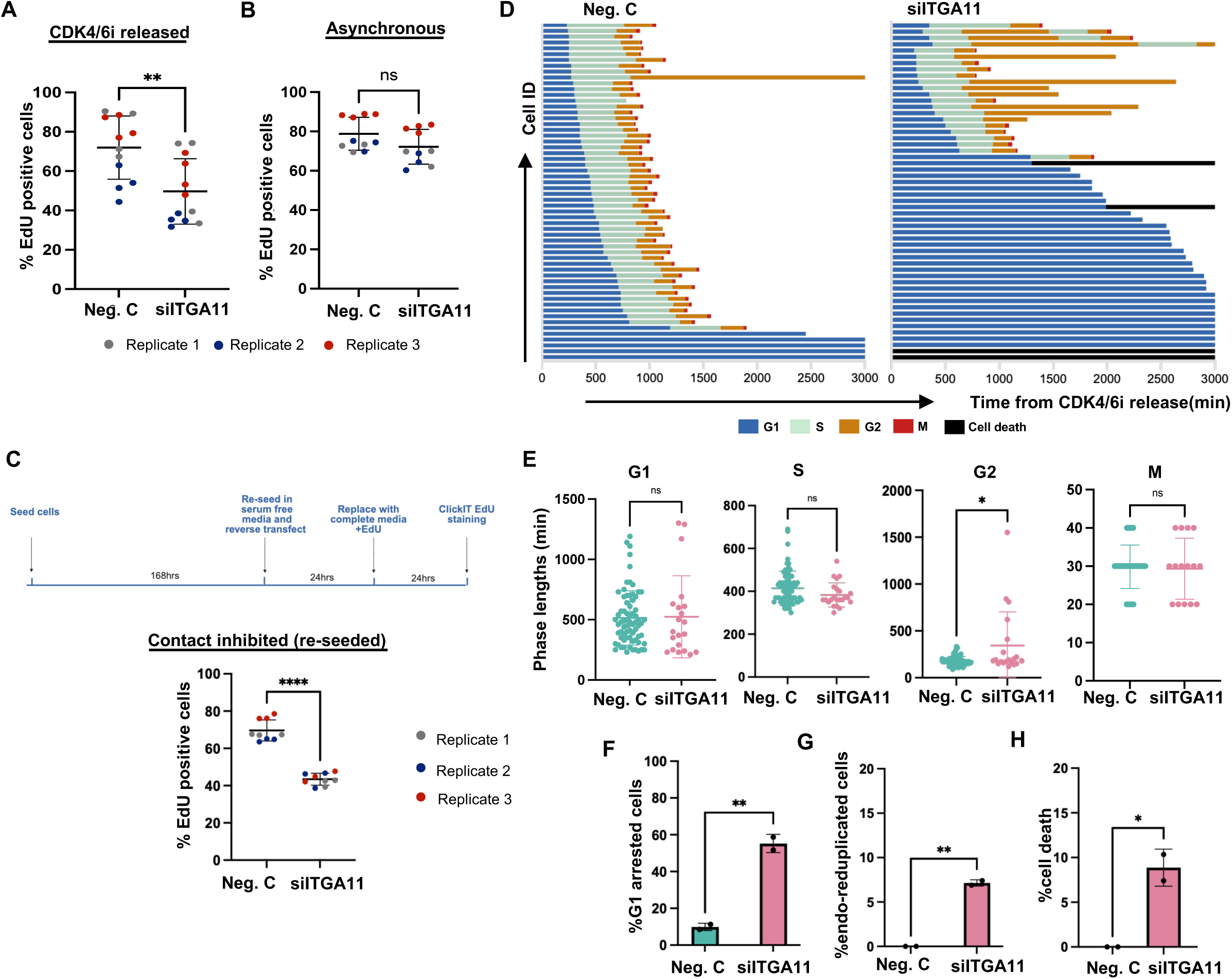
*ITGA11* promotes cell-cycle re-entry following quiescence. Quantification of EdU-positive cells following transfection with Neg. C or si*ITGA11* in (A) hTert-RPE1 cells released from CDK4/6i-induced quiescence and (B) asynchronously proliferating cells. Approximately 20% of Neg. C cells in B) are EdU negative, representing spontaneously quiescent cells^2^. Data are presented as mean ± SD from n=3 biological replicates, shown in grey, red and blue, respectively. Unpaired student’s t-test performed for statistical test, **P < 0.01, ns: not significant. C) (Top) Schematic of the experimental design used to assess cell-cycle re-entry following release from contact inhibition. Cells were transfected with Neg. C or si*ITGA11* immediately after replating in serum-free medium for 24hrs and then serum was added to release cells into cycle, and EdU incorporation was measured over the subsequent 24hrs. (Bottom) Quantification of EdU-positive cells following transfection with Neg. C or siITGA11 and release into the cell cycle. Data are presented as mean ± SD from three biological replicates, shown in grey, red and blue, respectively. Unpaired student’s t-test performed for statistical test, ****P < 0.0001. D) Representative single-cell cell-cycle traces of hTert-RPE1 mRuby-PCNA cells released from CDK4/6i arrest at t=0 mins. Black bars indicate cell death. Representative of n=2 biological replicates. E) Quantification of G1, S, G2, and M phase durations following release from CDK4/6i-induced quiescence. Data are shown as mean ± SD from three fields of view and are representative of n=2 biological replicates. Unpaired student’s t-test performed for statistical significance test, *P<0.05, ns: not significant. F) Percentage of cells arrested in G0/G1 following release from CDK4/6i in hTert-RPE1 mRuby-PCNA cells transfected with either Neg. C or si*ITGA11*. Unpaired student’s t-test performed to test for significance, **P < 0.01. G) Percentage of cells undergoing endo-reduplication in hTert-RPE1 mRuby-PCNA cells released from CDK4/6i and transfected with either Neg. C or si*ITGA11*. Unpaired student’s t-test performed to test for significance, **P < 0.01. H) Percentage of cells undergoing cell death in hTert-RPE1 mRuby-PCNA cells released from CDK4/6i and transfected with either Neg. C or si*ITGA11*. Unpaired student’s t-test performed to test for significance, *P < 0.05.

A similar cell cycle re-entry defect was observed in cells returning to proliferation from contact-inhibition (Fig. 3C; Supplementary Fig. 4B). In contrast to cells released from CDK4/6i-induced quiescence, *ITGA11*-depleted cells 48hrs post release from contact inhibition did not exhibit increased cell clustering compared to control-depleted cells (Supplementary Fig. 4C). This difference may reflect the disruption of cell-cell contacts during the re-seeding step required to release cells from contact inhibition.

Given the established role of *ITGA11* as a collagen-binding integrin that promotes cell-matrix adhesion, we next asked whether loss of *ITGA11* alters focal adhesion organization during cell cycle re-entry. To examine whether *ITGA11* influences adhesion dynamics, we quantified focal adhesion morphology in cells released from CDK4/6i-induced quiescence. *ITGA11* depleted cells displayed significantly smaller total area of pFAK-positive focal adhesions (average ∼30 µm²) compared with negative-control cells (∼40 µm²) (Supplementary Fig. 4D), consistent with impaired integrin-mediated adhesion maturation.

To analyse which cell cycle phases were perturbed by *ITGA11* loss, we assessed cell-cycle progression using live-cell imaging of hTert-RPE1 mRuby-PCNA cells^35^. In asynchronously cycling cells, *ITGA11* depletion did not alter the duration of G1, S, or G2 phases (Supplementary Fig. 5A-B), reinforcing that *ITGA11* is not required for continuous proliferation. However, we did observe increased cell death in *ITGA11*-depleted cells (Supplementary Fig. 5C). In contrast, cells released from two-day of CDK4/6 inhibition exhibited an increased proportion of cells not exiting quiescence and remaining in G1, consistent with fixed cell data (Fig. 3D-F). We also observed a prolonged G2 phase following *ITGA11* knockdown (Fig. 3E). Finally, we observed a higher frequency of endo-reduplication, where cells skip mitosis and re-enter S-phase, in *ITGA11*-depleted cells releases from CDK4/6i arrest compared to control depleted cells (Fig. 3G-H). Together, these data indicate defective completion of the first cell cycle after quiescence exit when *ITGA11* is depleted.

These findings indicate that *ITGA11* is dispensable for continuous proliferation, consistent with its low level of expression in these cells but becomes critical for efficient re-entry into the cell cycle after quiescence.

### *ITGA11* regulates cell-cycle re-entry through p27 stability

To elucidate the mechanism by which *ITGA11* loss impairs cell-cycle re-entry, we examined the expression of cell cycle proteins after *ITGA11* depletion. Notably, *ITGA11* depletion, either by siRNA or doxycycline-inducible shRNA, resulted in a marked increase in the protein levels of the CDK inhibitor protein, p27, in cells released from quiescence (Fig. 4A, Supplementary Fig. 6A-C) but not in asynchronously proliferating hTert-RPE1 mRuby-PCNA cells (Fig. 4B, Supplementary Fig. 6D). Although Cyclin D1 protein levels were also increased in *ITGA11*-depleted cells following release from CDK4/6i-induced quiescence (Supplementary Fig. 6E), these cells nevertheless failed to efficiently re-enter the cell cycle, consistent with elevated p27 overriding the pro-proliferative effects of Cyclin D1^36^.

**Figure 4:**
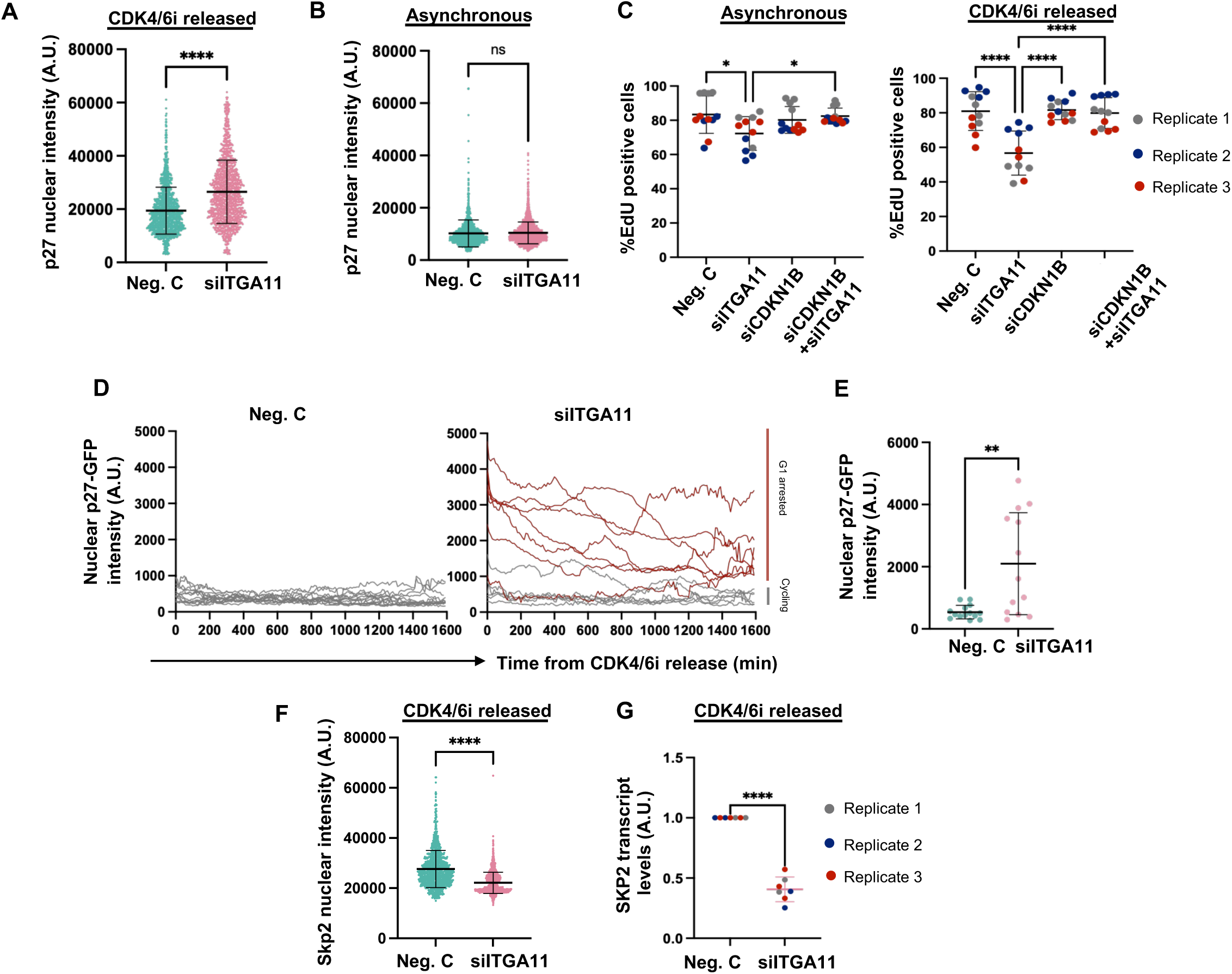
*ITGA11* regulates cell cycle re-entry through p27 stabilization. A) Quantification of nuclear p27 fluorescence intensity in CDK4/6i-released hTert-RPE1 mRuby-PCNA cells transfected with Neg. C or si*ITGA11*. Data are shown as mean ± SD and is representative of n = 3 biological repeats. Unpaired student’s t-test performed for comparison, ****P < 0.0001. B) Quantification of nuclear p27 fluorescence intensity in asynchronous (hTert-RPE1 mRuby-PCNA cells transfected with Neg. C or si*ITGA11*. Data are shown as mean ± SD and is representative of n = 3 biological repeats. Unpaired student’s t-test performed for comparison, ns:not significant. C) Percentage of cells undergoing cell cycle entry was assessed by EdU incorporation in asynchronously growing (left) and CDK4/6i-released (right) hTert-RPE1 mRuby-PCNA cells. Cells were transfected individually with Neg. C, si*ITGA11*, si*CDKN1B*, or a combination of si*ITGA11* and si*CDKN1B* for 48hrs, prior to the addition of EdU for 24hrs. Data are shown as mean ± SD (n = 3 biological repeats). One-way ANOVA statistical test performed for multiple comparisons, *P<0.05, ****P < 0.0001. D) Representative single-cell traces of nuclear p27-GFP intensity following release from CDK4/6i-induced quiescence. Traces are aligned to the time of release. Grey lines indicate cycling cells, and red lines indicate G0/G1-arrested cells. E) Quantification of nuclear p27-GFP intensity immediately after release (0 mins) from CDK4/6i-induced quiescence. Data are shown as mean ± SD and is representative of n = 2 biological repeats. Unpaired student’s t-test to test for significance, **P < 0.01. F) Quantification of nuclear Skp2 fluorescence intensity following release from CDK4/6i-induced quiescence in cells transfected with Neg. C or si*ITGA11*. Data are shown as mean ± SD and is representative of n = 3 biological repeats. Unpaired student’s t-test to test for significance, ****P < 0.0001. G) *SKP2* mRNA expression was measured by SYBR Green qPCR in cells transfected with either Neg. C or si*ITGA11* following 24hrs release from CDK4/6i-induced arrest. Data are shown as mean ± SD (n = 3 biological repeats). Unpaired student’s t-test performed for comparison, ****P < 0.0001.

Based on these results, we hypothesised that elevated p27 may be inhibiting cell cycle re-entry from quiescence in *ITGA11*-depleted cells. Simultaneous knockdown of p27 (encoded by *CDKN1B*, Supplementary Fig. 6F) and *ITGA11* in CDK4/6i-released cells fully rescued the cell-cycle arrest observed upon *ITGA11* depletion alone (Fig. 4C). These data suggest that *ITGA11* normally acts to reduce p27 protein levels to promote cell cycle entry.

We next investigated when *ITGA11* loss leads to elevated p27 protein levels. Live-cell imaging of hTert-RPE1 mRuby-PCNA p27-GFP cells (where p27 is labelled at the endogenous locus with GFP^37^); revealed that *ITGA11*-depleted cells displayed elevated nuclear p27-GFP levels following release from CDK4/6i-induced quiescence compared with negative control cells (Fig. 4D). Quantification of nuclear p27-GFP intensity demonstrated that p27 levels were already significantly elevated at the time of release from quiescence (time 0 mins) in *ITGA11*-depleted cells (Fig. 4E), indicating that p27 accumulates during quiescence in the absence of *ITGA11*, before cell-cycle re-entry.

p27 protein is degraded after ubiquitination via the SCF^SKP2^ E3 ubiquitin ligase complex^38, 39^. Therefore, we examined whether *ITGA11* influences Skp2 abundance. Both Skp2 protein (Fig. 4F, Supplementary Fig. 6G) and *SKP2* mRNA (Fig. 4G) were significantly decreased in *ITGA11*-depleted cells released from CDK4/6i-induced quiescence.

These findings indicate that *ITGA11* positively regulates *SKP2* expression, thereby enabling efficient p27 turnover during cell cycle re-entry.

### *ITGA11* regulates cell-cycle re-entry through YAP-dependent control of Skp2 and p27

Mechano-transduction pathways provide a compelling mechanistic link between integrin signalling and nuclear control of proliferation. The Hippo pathway effector YAP is a mechano-responsive transcriptional co-activator that integrates ECM stiffness^40, 41^, cytoskeletal tension, and integrin engagement to regulate G1-S progression^32, 33^.

YAP promotes transcription of *SKP2*^42^, and modulates the Cyclin D1/p27 ratio in response to ECM cues^36^. Because p27 stability is a critical determinant of quiescence maintenance and exit, perturbations in ECM-integrin-YAP signalling could profoundly influence the decision to re-enter the cell cycle.

We first assessed YAP abundance following *ITGA11* depletion. Cells released from CDK4/6i-induced quiescence exhibited reduced total YAP protein levels upon *ITGA11* knockdown, compared to control depleted cells (Fig. 5A). Live-cell imaging of overexpressed EGFP-YAP1 showed a reduction in nuclear to cytoplasmic YAP1 following *ITGA11* loss in CDK4/6i released cells (Fig. 5B-C) suggesting impaired nuclear translocation and transcriptional activity, in addition to reduced YAP1 levels shown by western blotting (Fig. 5A). Together, these findings demonstrate that *ITGA11* is required to maintain YAP1 abundance and nuclear localization during cell-cycle re-entry, consistent with reduced YAP1 transcriptional activity contributing to the decreased *SKP2* expression and p27 accumulation observed following *ITGA11* depletion.

**Figure 5:**
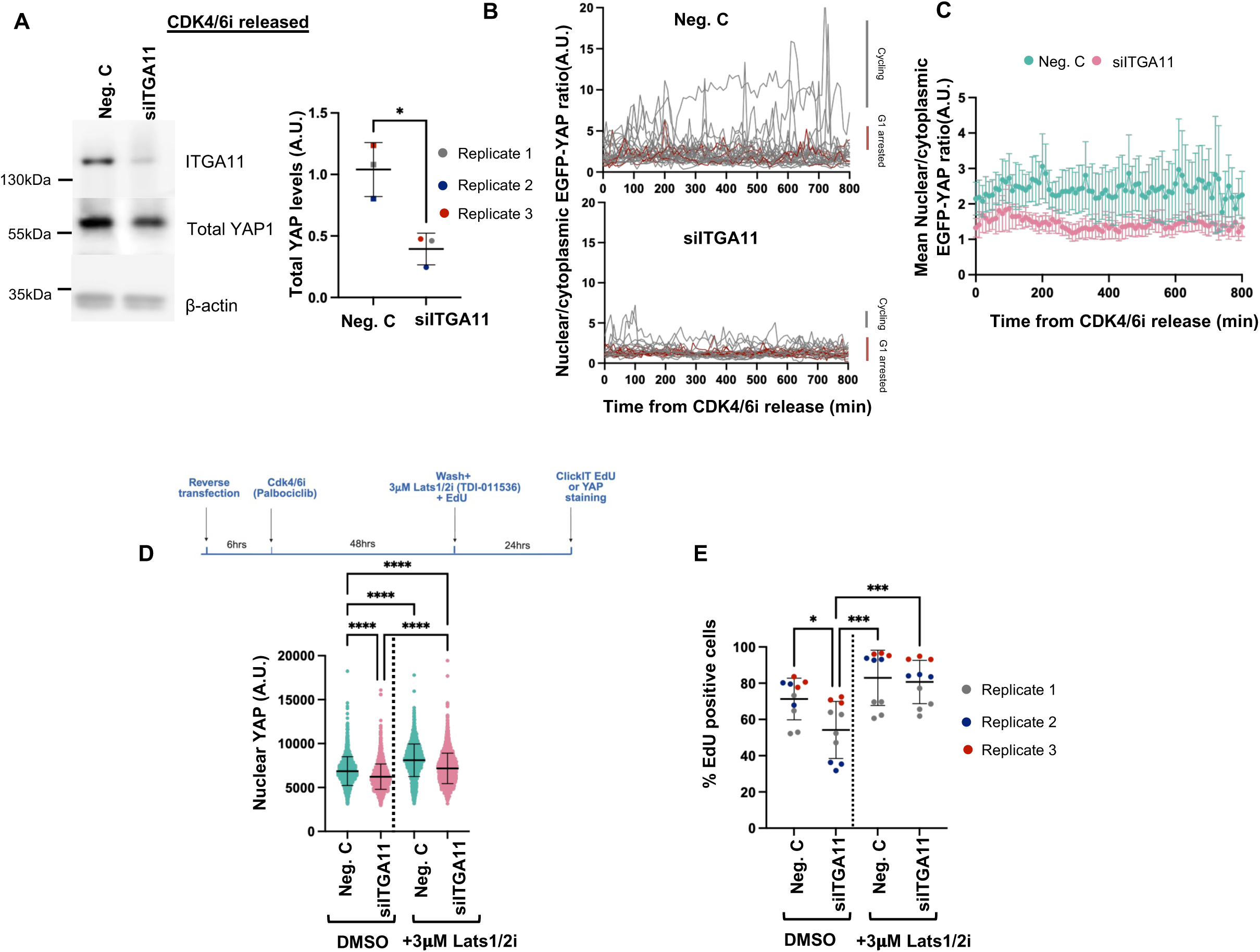
ITGA11 regulates cell-cycle re-entry through YAP-dependent control of SKP2 and p27. A) Western blot of whole cell lysates showing the levels of total YAP in Neg.C and si*ITGA11* cells released from CDK4/6i. Quantification of YAP normalized to β-actin. Data are shown as mean ± SD (n = 3 biological repeats). Unpaired student’s t-test performed for comparison, *P<0.05. B) Single cell traces of EGFP-YAP1 nuclear to cytoplasmic ratio after hTert-RPE1 mRuby-PCNA EGFP-YAP1 cells are released from CDK4/6 inhibitor-induced arrest. Data shown are representative of n=2 biological repeats. Grey lines indicate cycling cells, and red lines indicate G1-arrested cells. C) Mean EGFP-YAP1 nuclear to cytoplasmic ratio after hTert-RPE1 mRuby-PCNA EGFP-YAP1 cells are released from CDK4/6 inhibitor-induced arrest. Data are presented as mean ± 95% CI (n = 29 and 25 cells for Neg.C, and n = 23 and 20 cells for si*ITGA11*). D) (Top) Schematic representation of the experimental design used to assess cell-cycle re-entry in hTert-RPE1 mRuby-PCNA cells released from CDK4/6i-induced quiescence and treated with DMSO or 3μM LATS1/2 inhibitor (TDI-011536) following transfection with the indicated siRNAs. (Bottom) Quantification of nuclear YAP fluorescence intensity. Due to extensive cell clustering observed in si*ITGA11* cells released from CDK4/6i, nuclear YAP intensity was used as a measure of YAP activity (instead of nuclear: cytoplasmic ratio)^65^. Data shown are representative of n=3 biological repeats. One-way ANOVA statistical test performed for multiple comparisons, ****P < 0.0001. E) Percentage of EdU-positive cells in hTert-RPE1 mRuby-PCNA released from CDK4/6i induced arrest were treated with 3ìM LATS1/2 inhibitor (TDI-011536) or DMSO in different siRNA conditions. Data are shown as mean ± SD (n = 3 biological repeats). One-way ANOVA statistical test performed for multiple comparisons, *P<0.05, ***P < 0.001.

To determine whether reduced YAP transcriptional activity accompanies these changes, we quantified transcript levels of canonical YAP target genes, *CTGF* and *CYR61* in cells released from CDK4/6i-induced quiescence. Both transcripts were significantly decreased in *ITGA11*-depleted cells (Supplementary Fig. 7A), consistent with decreased *SKP2* transcription (Fig. 4G) and diminished YAP1-dependent transcription.

Given that YAP nuclear accumulation is restrained by LATS1/2-mediated phosphorylation^43^, we asked whether pharmacological inhibition of LATS1/2 could enhance nuclear YAP and rescue the *ITGA11*-dependent cell-cycle defect. Treatment of *ITGA11*-depleted cells with a LATS1/2 inhibitor following release from CDK4/6i-induced quiescence restored nuclear YAP localization (Fig. 5D) and rescued the cell cycle re-entry defect after *ITGA11* depletion (Fig. 5E). To assess the specificity and functional activity of the inhibitor, we examined the expression of the canonical YAP target genes *CTGF* and *CYR61*. Expression of both genes was restored following LATS1/2 inhibitor treatment in cells released from quiescence (Supplementary Fig. 7B). Consistent with these findings, LATS1/2 inhibition reduced YAP phosphorylation at Ser127 and increased YAP protein levels (Supplementary Fig. 7C). The rescue of cell-cycle re-entry was accompanied by increased Skp2 expression and reduced p27 levels (Supplementary Fig. 7D, E).

Together, these findings indicate that restoring YAP nuclear activity is sufficient to bypass the cell-cycle arrest induced by *ITGA11* depletion.

Thus, *ITGA11*-dependent regulation of YAP nuclear activity represents a key mechanism controlling cell-cycle re-entry following quiescence.

## Discussion

While the molecular mechanisms governing entry into quiescence have been extensively characterized, the pathways that enable quiescent cells to retain proliferative competence remain incompletely understood. Using quantitative proteomics to compare quiescent, senescent and proliferating cells, followed by functional image-based screening, we identified a set of quiescence-enriched proteins that regulate the transition from quiescence to proliferation. Among these, we identify the collagen I receptor integrin α11 (*ITGA11)* as a previously unrecognized regulator of quiescence exit. *ITGA11* was consistently induced across multiple models of reversible quiescence and in different cell types, while remaining low in senescent cells, identifying it as a functional marker of reversible cell-cycle arrest rather than a general marker of growth arrest. Unlike many cell-cycle regulators that broadly affect proliferation, *ITGA11* was specifically required for re-entry from quiescence while being largely dispensable for asynchronously proliferating cells. These findings indicate that quiescent cells acquire specialized ECM signalling mechanisms that prepare them for subsequent proliferation. Mechanistically, our data place *ITGA11* upstream of a YAP-Skp2-p27 signalling axis that promotes p27 turnover during quiescence exit (Fig. 6). This work identifies *ITGA11* as a functional quiescence signature protein and suggests that ECM sensing is an integral component of the molecular program that licenses quiescent cells to regain proliferative competence. Our findings are consistent with previous transcriptomic studies reporting enrichment of ECM- and cell adhesion-associated gene signatures in quiescent cells^2, 44^,but extend these observations by demonstrating that *ITGA11* is not merely a marker of quiescence but is functionally required for efficient quiescence exit.

**Figure 6:**
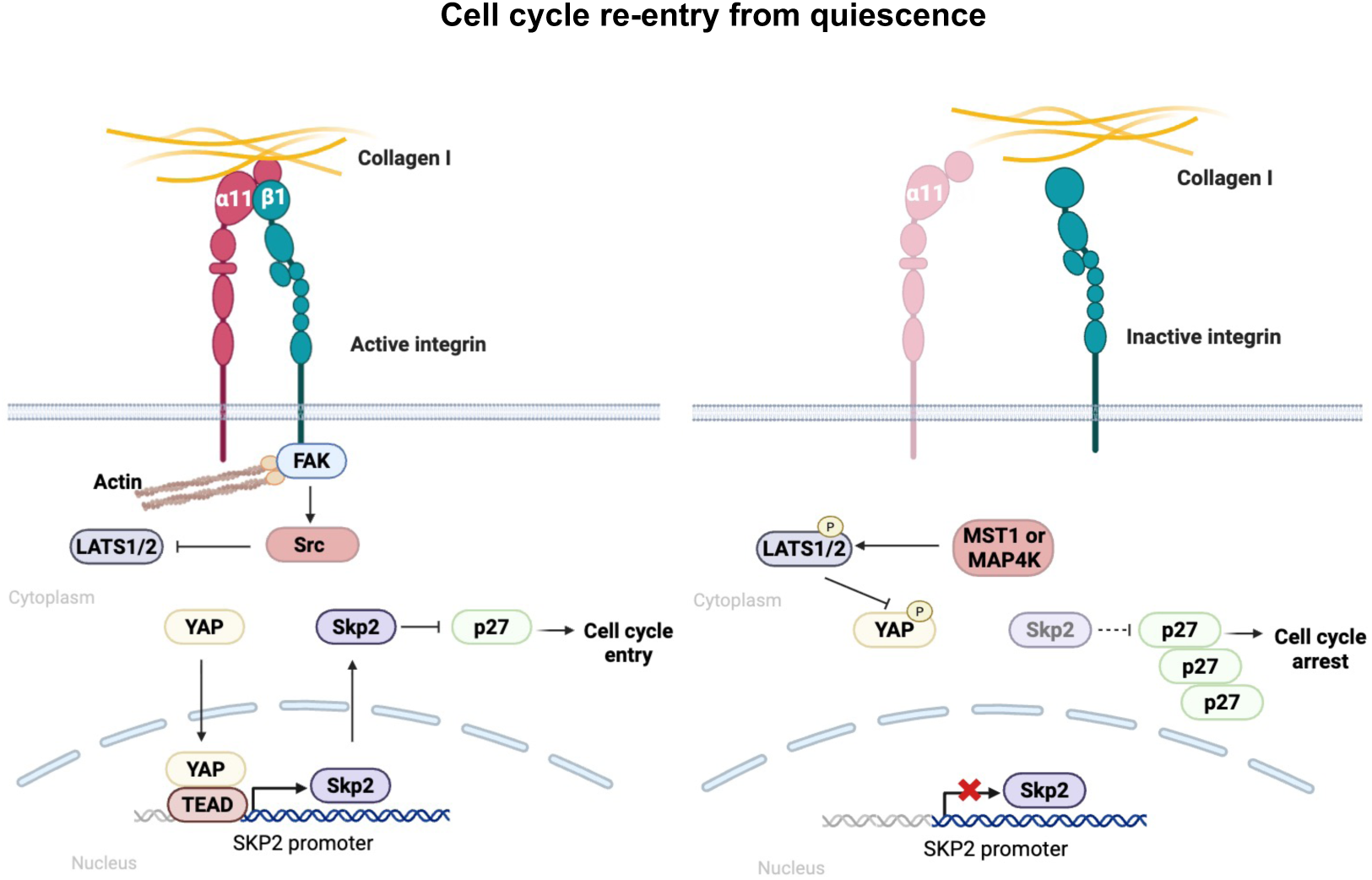
Proposed model for integrin alpha11 (α11)-mediated regulation of cell cycle re-entry from quiescence through YAP signalling. (Left) In cells re-entering the cell cycle from quiescence, collagen I-bound ITGA11 activates FAK and Src signalling, promoting YAP nuclear localization and SKP2 transcription, which facilitates p27 degradation and supports cell-cycle entry. (Right) In the absence of ITGA11, impaired integrin signalling activates the MST1/2-LATS1/2 pathway, leading to increased YAP phosphorylation and cytoplasmic sequestration. Reduced nuclear YAP decreases SKP2 expression, resulting in p27 stabilization and failure to re-enter the cell cycle.

Integrin signalling has established roles in cell proliferation through FAK, SRC, MAPK and PI3K signalling pathways^45^. In contrast, *ITGA11* has primarily been studied as a fibroblast-specific collagen receptor involved in extracellular matrix remodelling and fibrosis^46, 47^. However, *ITGA11* expression extends beyond fibroblasts, with high expression observed across mesenchymal and smooth muscle lineages as well as other specialized cell populations^48^. This broader expression pattern is consistent with accumulating evidence that matrix composition and mechanical cues regulate stem cell activation, tissue regeneration, and fibroblast behaviour through integrin-dependent mechano-transduction pathways^40, 49, 50^. More broadly, our findings provide a potential mechanistic explanation for the reduced body size reported in *ITGA11-* deficient mice, suggesting that impaired *ITGA11*-dependent quiescence exit may limit proliferative expansion during development. This model parallels observations in *SKP2*-knockout mice, in which defective cell cycle re-entry contributes to reduced organismal growth^51^, raising the possibility that efficient exit from quiescence is an important determinant of normal tissue growth and homeostasis.

An intriguing observation from this study is that *ITGA11* depletion reduced not only the nuclear localization of YAP but also its overall protein abundance. YAP activity is regulated at multiple levels, including transcription, protein stability, phosphorylation-dependent degradation, and nucleo-cytoplasmic shuttling^52^. Mechanical inputs such as integrin engagement, focal adhesion signalling, actomyosin contractility, and extracellular matrix stiffness can influence both YAP stability and nuclear accumulation through regulation of the Hippo pathway and associated signalling networks^40, 53^. While our rescue experiments place YAP downstream of *ITGA11*, the precise mechanism by which *ITGA11* controls YAP abundance remains unclear. One possibility is that *ITGA11*-dependent collagen sensing promotes cytoskeletal tension and suppresses Hippo pathway activity by preventing activation of core Hippo kinase MST1, thereby stabilizing YAP and facilitating its nuclear accumulation. Alternatively, *ITGA11* may influence YAP expression or turnover through FAK-SRC-dependent signalling pathways. Future studies examining YAP transcription, phosphorylation status, and protein stability following *ITGA11* depletion will be required to distinguish between these possibilities.

An additional finding of this study is the selective enrichment of *ITGA11* in quiescent, but not senescent, cells. Distinguishing reversible quiescence from irreversible senescence remains a major challenge because of the lack of robust markers that reliably discriminate these two non-proliferative states. The preferential expression of *ITGA11* in quiescent cells, together with its localization at the cell surface, raises the possibility that *ITGA11* could serve as a biomarker for identifying and isolating reversibly arrested cells. The lower proportion of *ITGA11*-positive cells detected by flow cytometry relative to immunofluorescence likely reflects methodological differences, as flow cytometry quantifies only cell surface-localized *ITGA11*, whereas immunofluorescence detects both intracellular and surface-associated pools of the protein. Future studies will be required to determine whether integrin α11 can be exploited to prospectively isolate quiescent cells from heterogeneous tissues and to distinguish quiescent from senescent cells in physiological and pathological contexts.

Finally, these findings may have broader implications beyond cell-cycle control. *ITGA11* is highly expressed in activated fibroblasts and cancer-associated fibroblasts, where it contributes to extracellular matrix remodelling and tissue stiffness^54, 55^. Our results raise the possibility that *ITGA11* regulates transitions between quiescent and activated fibroblast states through YAP-dependent signalling, with potential relevance to fibrosis, wound repair, and tumour progression.

Collectively, our findings establish an ITGA11-YAP-Skp2-p27 signalling axis that couples extracellular matrix sensing to the regulation of cell-cycle re-entry. We propose that *ITGA11* acts as a collagen-responsive integrin that primes quiescent cells for proliferative competence by sustaining YAP activity, maintaining Skp2 expression, and facilitating p27 turnover upon exit from quiescence. These findings provide a mechanistic framework for how extracellular matrix-derived signals are integrated with intracellular cell-cycle control pathways to govern the transition from reversible cell-cycle arrest to proliferation.

## Methods

### Cell culture

hTert-RPE1 mRuby-PCNA cells were obtained from Joerg Mansfeld (ICR, London)^35^. hTert-RPE1 mRuby-PCNA p27-GFP in which the endogenous *CDKN1B* gene was tagged at the C-terminus using CRISPR-mediated gene tagging, were generated as described previously^37^.

All cell lines were cultured in DMEM (Gibco) supplemented with 10% fetal bovine serum (FBS) and 1% penicillin–streptomycin at 37 °C in a humidified incubator with 5% CO₂. The cell lines were regularly tested for mycoplasma.

### Cell line generation

*Generation of hTert-RPE1 mRuby-PCNA mH2.B-mTurquoise EGFP-YAP1 cell line* hTert-RPE1 mRuby-PCNA were first tagged with *mH2.B-mTurquoise* using CSII-EF lentiviral vector ^56^. HEK293T cells were transfected with 3µg of this plasmid together with packaging and envelope plasmids (7.5µg pMDLg/pRRE, 7.5µg pRSV/REV, and 5µg pMD2.G) using Lipofectamine 2000 (Thermo Fisher Scientific, 11668019) according to the manufacturer’s protocol. 48hrs post-transfection, viral supernatant was collected and filtered through a 0.45µm filter. hTert-RPE1 mRuby-PCNA cells were transduced with 300µl of diluted virus in the presence of polybrene (final concentration 1µg/ml). Cells were subsequently single-cell sorted using a BD FACSAria cell sorter into 96-well plates containing 100µl of conditioned media based on mTurquoise expression and expanded in complete medium.

These cells were forward-transfected with 3µg of pEGFP-C3-hYAP1 plasmid^57^ using Lipofectamine 2000 (Thermo Fisher Scientific, 11668019) following the manufacturer’s instructions. After 24hrs, GFP-positive cells were single-cell sorted using a BD FACSAria cell sorter into 96-well plates and maintained in complete medium containing 1mg/ml G418 for selection. After selection, cells were maintained in 500µg/ml of G418.

*Generation of hTert-RPE1 mRuby-PCNA doxycycline-inducible shITGA11 cell lines* <u>shRNA cloning.</u> To generate shRNAs targeting *ITGA11*, the following oligonucleotides were cloned into EcoRI- and AgeI-digested Tet-pLKO.1-Neo ^58^ vector:

*Control* Forward:5’CCGGCAAGCTGACCCTGAAGTTCATCTCGAGATGAACTTCAGGGTCGCTTGTTTTTG 3’

Reverse:5’AATTCAAAAACAAGCTGACCCTGAAGTTCATCTCGAGATGAACTTCAGGGTCAGCTTG 3’

*shRNA1* Forward:5′CCGGGGAACTGCACCAAACTCAACCCTCGAGGGTTGAGTTTGGTGCAGTTCCTTTTTG-3′ Reverse:5′AATTCAAAAAGGAACTGCACCAAACTCAACCCTCGAGGGTTGAGTTTGGTGCAGTTCC-3′

*shRNA2* Forward:5′CCGGGGAGAAGGTGATCCAGCAAAGCTCGAGCTTTGCTGGATCACCTTCTCCTTTTTG-3′ Reverse:5′AATTCAAAAAGGAGAAGGTGATCCAGCAAAGCTCGAGCTTTGCTGGATCACCTTCTCCC-3′

*shRNA3* Forward:5′CCGGGCACGACATCAGTGGCAATAACTCGAGTTATTGCCACTGATGTCGTGCTTTTTG-3′ Reverse:5′AATTCAAAAAGCACGACATCAGTGGCAATAACTCGAGTTATTGCCACTGATGTCGTGC-3′

*shRNA4* Forward:5′CCGGAGTTACCAGAATGCCCGATTTCTCGAGAAATCGGGCATTCTGGTAACTTTTTTG-3′ Reverse:5′AATTCAAAAAAGTTACCAGAATGCCCGATTTCTCGAGAAATCGGGCATTCTGGTAACT-3′

Lentiviral particles were produced as described above. hTert-RPE1 mRuby-PCNA cells were transduced with viral supernatant and selected with 1mg/ml G418. To induce knockdown of *ITGA11*, doxycycline (1µg/ml; Sigma-Aldrich, D9891) was added for 24hrs. Knockdown efficiency was validated by immunofluorescence.

### In vitro quiescent models

#### CDK4/6 inhibitor induced

hTert-RPE1 cells were treated with the CDK4/6 inhibitor Palbociclib (Selleckchem, S1116) at a final concentration of 1µM for 48hrs^26^. Cells were then washed three times with PBS and released back into the cell cycle by incubation in complete growth medium for 24hrs.

#### Contact inhibition

hTert-RPE1 cells were arrested in G0/G1 by contact inhibition by allowing cultures to reach full confluence and maintaining them for 7 days in a 6-well plate. Cells were subsequently replated at low density (1000 cells/well) in serum-free DMEM for 24hrs and transfected with siRNAs.

### Fluorescence-activated cell sorting

3×10^6^ hTert-RPE1 mRuby-PCNA p21-GFP cells were seeded and allowed to attach for 24hrs. Cells were then treated with 50µM Etoposide for 3 days to induce DNA damage. Following treatment, the drug was washed out, and cells were cultured for an additional 3 days to promote senescence. Senescence was confirmed by increased p21 expression and β-galactosidase positivity.

Asynchronously growing and etoposide-treated hTert-RPE1 mRuby-PCNA p21-GFP cells were sorted using a BD FACSAria cell sorter with BD FACSDiva software (v9.4). Cells were separated based on mRuby and p21-GFP fluorescence into three populations: cycling G1 cells (p21low PCNAlow), G0 cells (p21high PCNAlow), and senescent cells (p21high). For each population 1×10^6^ cells were sorted per sort and three biological repeats were performed. After sorting, cells were frozen down in complete media with 10% DMSO before being processed for proteomics.

### Sample preparation and data acquisition, analysis for Mass Spectrometry

Cells were lysed with 2% SDS in PBS supplemented with protease and phosphatase inhibitors. Cell lysates were homogenised by bath sonication and nucleic acids were digested with Benzonase for 30min at 37 °C. Proteins were precipitated with acetone and digested into peptides with trypsin (1:100 enzyme to protein ratio) for 16 h at 37 °C, followed by a second round of digestion for 4hrs at 37 °C. Next, peptides were acidified with formic acid (final concentration of 3%) and desalted with Waters SepPak C18 columns. Peptides were eluted using 80% acetonitrile in 0.5% formic acid and dried in a vacuum concentrator at 30 °C. Next, each sample was dissolved in 100 mM TEAB, mixed with 0.25mg of TMT label and incubated for 1hr at room temperature. Labelling reaction was quenched by adding 5% hydroxylamine to each sample and incubating for 15 min at 37 °C as recommended by manufacturer’s protocol. Next, all samples were pooled together and dried before C18 desalting using Waters Sep-Pak C18 columns. To achieve a deep coverage of the phosphoproteome, peptides were fractionated using high-pH reversed-phase chromatography. Samples were resuspended in 10 mM ammonium formate pH 9.0 and fractionated using an Ultimate 3000 HPLC (ThermoFisher Scientific) equipped with a XBridge Peptide BEH C18 130 Å 3.5μm 4.6 × 150 mm column (Waters). Peptides were eluted at 1 ml/min using a constant 10 mM ammonium formate pH 9.0 and a multistep gradient from 2% to 50% acetonitrile over 10min, 50% to 80% gradient for a further 1min, followed by an 80% column wash and re-equilibration. 16 concatenated fractions were collected. Peptides were eluted, dried and stored at −20 °C until measurement by LC-MS.

Peptides were analysed using the Orbitrap Fusion Lumos Tribrid mass spectrometer. coupled online, to an Ultimate 3000 HPLC (Dionex, ThermoFisher Scientific, UK). Peptides were separated on a 50 cm (2-µm particle size) EASY-Spray column (Thermo Scientific, UK). Mobile phase A consisted of 0.1% formic acid in LC-MS grade water and mobile phase B consisted of 80% acetonitrile and 0.1% formic acid. Peptides were loaded onto the column at a flow rate of 0.3 μl min−1 and eluted at a flow rate of 0.25 μl min−1 according to the following gradient: 2–40% mobile phase B in 120 min and then to 95% in 11 min. Mobile phase B was retained at 95% for 5 min and returned back to 2% a minute after until the end of the run (160 min). Eluted peptides were then ionised using an EASY-Spray source (Thermo Scientific, UK), which was operated at 50 °C.

Ions were mass analysed using Orbitrap MS1 scans recorded at 120,000 resolution (scan range 380–1500 m/z) with an ion target of 4.0e5, and injection time of 50 ms. MS2 CID fragmentation was performed in the linear ion trap in turbo scan mode, with an ion target of 2.0E4 and a normalised collision energy of 35, Q activation parameter at 0.25 and CID activation time at 10 ms. MS3 scans were performed in the Orbitrap, with the number SPS precursors set to 5. The isolation window was set to 2 Th, and the resolution at 50,000 with scan range 100–150 m/z. MS3 HCD fragmentation was performed with normalised collision energy of 65.

Raw data were processed using MaxQuant (version 1.6.2.6) with the default settings for a TMT 10-plex experiment and included corrections for isotopic contamination according to manufacturer’s recommendations. A maximum of 2 missed cleavages were allowed and the enzyme parameter was set to ‘Trypsin/P’. Carbamidomethyl (C) was added as a fixed modification while Oxidation (M), Acetyl (Protein N-term), Deamidation (NQ), Gln-> pyroGlu and Phospho (STY) were added as variable modifications with a maximum number of 5 modifications per peptide. Note that Phospho (STY) was considered as a variable modification because the dataset included samples processed in parallel for phosphoproteomics. Phosphorylation data were not included in this study. Database search was done against the human Swiss-Prot database, accessed in June 2018. Protein intensities were divided to total intensity per channel and multiplied by 1 million to obtain parts-per-million (ppm) abundances. Pairwise t-tests were performed to assess differential protein expression.

### siRNA screens

A custom library of ONTarget siRNA Pools (ThermoFisher) targeting 129 genes encoding proteins (including controls) as high confidence hits in our proteomics analyses were ordered. ONTarget non-targeting control pool siRNA was used as a negative control. ONTargetPlus pool siRNA targeting *PLK1* was used as a positive control as it kills hTert-RPE1 cells and serves as a visual check that the screen is working. Screen controls included ONTargetPlus pool siRNAs targeting *TP53*, as this promotes faster cell cycle entry and a reduced fraction of EdU negative cells^1^, and *CCND1*, which is essential for proliferation in hTert-RPE1 cells ^35^and so its depletion promotes a high fraction of EdU negative cells. siRNAs were diluted and dispensed onto 384 well Phenoplates (Revvity) using an Echo acoustic liquid handler −160nl of a 5μM stock per 384well. Plates were stored at −80°C until use. All screens were performed in technical replicate. For the screens, plates were thawed and siRNAs resuspended in 5μl OPTIMEM and briefly centrifuged. Lipofectamine RNAiMAX was diluted 40nl/5μl in OPTIMEM and 5μl added to each siRNA containing well. Plates were briefly centrifuged. hTert-RPE1 cells were added in 20μl of complete growth medium to each well at a concentration of 0.25×10^5^/ml (500 cells/well) and plates briefly centrifuged.

For the asynchronous screen, cells were transfected as above and incubated for 48hrs. 5μM EdU was added for the last 24hrs of this incubation.

For the two- and three-day CDK4/6 inhibitor arrest and release screen, cells were treated as above and after 6hrs, 1μM CDK4/6i was added and cells incubated for two- or three-days. After the incubation, cells were washed three times in PBS and released into full growth medium containing 5μM EdU.

At the end of the incubation, plates were fixed in 4% formaldehyde and processed for immunostaining (as below).

### siRNA transfection

Custom siRNA targeting CDKN1B CAAGUGGAAUUUCGAUUUUtt (Ambion Silencer Select s2837) and siITGA11 pools from siTOOLS Biotech were added at a final concentration of 5nM. Briefly, 7.5nl of Lipofectamine RNAiMax was added to 7.5nl 20 μM (stock concentration) siRNA in 10μl OptiMEM (Gibco 31985062) and then added to 750 cells/well resuspended in 20μl complete media. For CDK4/6i released condition, 6hrs after transfection cells were treated with 1μM CDK4/6i and incubated for 48hrs. Cells were washed and released into full growth medium containing 5μM EdU. For asynchronous, EdU was added 48hrs after transfection and incubated for 24hrs.

Contact inhibited cells were harvested and re-seeded at 1000 cells/well, reverse transfected with siRNAs and grown in serum-free media for 24hrs as described earlier. Cell cycle re-entry was induced by the addition of serum-containing DMEM for 24hrs along with 24hrs EdU treatment.

For western blotting, cells were reverse transfected with siRNA in 24 well plates (Greiner Bio-One, cat. No. 662165) in a total volume of 500μl. Briefly, 0.125μl of Lipofectamine RNAiMax was added to 0.125μl 20μM (stock concentration) siRNA in 150μl OptiMEM (Gibco 31985062) and then added to cells.

### Immunostaining

Cells were seeded in 96- or 384-well PhenoPlates (PerkinElmer, 6055302 and 6057302, respectively). EdU was added either 48hrs after transfection or following release from CDK4/6 inhibitor treatment to a final concentration of 10μM and incubated for 24hrs prior to fixation. Cells were fixed in 4% formaldehyde in PBS for 15 min at room temperature (RT), permeabilized with PBS containing 0.25% Triton X-100 for 10 min, and blocked for 1hr in blocking buffer (PBS supplemented with 2% BSA).

Primary antibodies were diluted in blocking buffer and incubated with cells overnight at 4 °C. After three washes in PBS, cells were incubated for 1hr at RT in the dark with appropriate secondary antibodies (Invitrogen).

For EdU detection, cells were incubated in click-reaction buffer containing 100 mM Tris-HCl (pH 7.5), 4 mM CuSO₄, 100 mM ascorbic acid, and 5μM sulfo-cyanine-5 azide for 40min at RT in the dark. Cells were then washed three times with PBS and counterstained with Hoechst 33582 (1μg/ml) for 8min, followed by three additional PBS washes. All washing steps were performed using an automated 50TS microplate washer (BioTek).

Plates were imaged using an Operetta CLS high-content imaging system (Revvity) equipped with either a 20× objective (NA 0.8) for EdU imaging or a 40× water objective (NA 1.1) for ITGA11 and pFAK imaging in the confocal mode.

Primary antibodies used for immunostaining included p27Kip1 (clone D37H1, CST 3688, 1:500), integrin α11 or ITGA11 (clone 203E1, Nanotools 0518-100, 1:200), YAP (clone 63.7, SC101199, 1:200), Skp2 (clone D3G5, CST 2652, 1:500), pFAK (Thermo Scientific 44-626G, 1:200), and vinculin (CST 13901, 1:500). Secondary antibodies were goat anti-mouse Alexa Fluor-488 (Invitrogen A11001, 1:1000), goat anti-mouse Alexa Fluor-647 (Invitrogen A21235, 1:1000), goat anti-rabbit Alexa Fluor-488 (Invitrogen A11008, 1:1000), and goat anti-rabbit Alexa Fluor-647 (Invitrogen A21245, 1:1000).

### Real time PCR

hTert-RPE1 mRuby-PCNA cells (1 × 10^5^ cells per condition) were collected from 6-well plates under asynchronous growth conditions, following 2 days of palbociclib treatment, 7 days of contact inhibition, or treatment with 50μM etoposide to induce senescence. Total RNA was extracted using the RNeasy kit according to the manufacturer’s instructions. cDNA was synthesized from 400ng of total RNA using LunaScript RT SuperMix.

Quantitative PCR (qPCR) was performed using SYBR Green master mix with forward and reverse primers at a final concentration of 500nM each. Reactions were prepared in a total volume of 20μL containing 10μL SYBR Green master mix, 1μL forward primer (10μM), 1μL reverse primer (10μM), 1μL cDNA, and 7μL nuclease-free water. Relative gene expression was determined using the ΔΔCt method following normalization to GAPDH housekeeping gene.

Primers used:

*ITGA11*

Forward: 5’ CAGCTCGCTGGAGAGATACG 3’

Reverse: 5’ TTACAGGACGTGTTCGCCTC 3’

*MAX*

Forward: 5’ AACGTAGGGACCACATCAAAG 3’

Reverse: 5’ AAGACAGGTGGGAGGGTTA 3’

*EZH2*

Forward: 5’ TTGTTGGCGGAAGCGTGTAAAATC 3’

Reverse: 5’ TCCCTAGTCCCGCGCAATGAGC 3’

*SKP2*

Forward: 5’ ATGCCCCAATCTTGTCCATCT 3’

Reverse: 5’ CACCGACTGAGTGATAGGTGT 3’

*CTGF*

Forward: 5’ CTGTGCAGCATGGACGTTCG 3’

Reverse: 5’ CATTTCCCGGGCAGCTTGAC 3’

*CYR61*

Forward: 5’ GGCTGGAATGCAACTTCGGC 3’

Reverse: 5’ ACAGAGGAATGCAGCCCACG 3’

*GAPDH*

Forward: 5’ GTCAACGGATTTGGTCGTATTG 3’

Reverse: 5’ TGTAGTTGAGGTCAATGAAGGG 3’

### Flow Cytometry

hTert-RPE1 mRuby-PCNA cells were seeded at 1,000 cells per well in 96-well plates and cultured either asynchronously or treated with CDK4/6 inhibitor Palbociclib (Selleckchem, S1116) for 48hrs or contact inhibited as before. Integrin α11 antibody (clone 203E1, Nanotools 0518-100) was diluted to a final concentration of 3μg/mL and conjugated to Zenon Alexa Fluor 488 according to the manufacturer’s instructions. Cells were detached using Accutase, resuspended in complete medium, and incubated with the ITGA11-Zenon 488 complex for 1hr in the dark with gentle shaking. Cells were then washed twice with 1X PBS, resuspended in 200ml of1XPBS, and analysed on a Symphony flow cytometer using a plate-based acquisition system. The data was analysed on FlowJo 10.10.0.

### Image analysis

Automated image analysis of images acquired on the Operetta CLS was performed using Harmony software (Revvity). Nuclei were segmented based on Hoechst intensity, and nuclei located at the edge of the field of view were excluded from analysis. Nuclear protein intensity was calculated as the mean intensity within the segmented nuclear region.

For quantification of integrin α11 cytoplasmic intensity, the cytoplasmic compartment was segmented based on vinculin intensity. Nuclear and cytoplasmic integrin α11 fluorescence intensities were quantified as the mean fluorescence intensity within their respective segmented regions.

Quiescent cells were identified based on the absence of EdU incorporation. A nuclear-to-cytoplasmic (N:C) ratio was calculated by generating a four-pixel wide ring surrounding each segmented nucleus to mark the cytoplasmic region. N:C ratios for individual cells were calculated and plotted using GraphPad Prism. Cells with EdU N:C ratios below 1.2 were classified as G0.

For the image-based screen, Z-scores were calculated for each siRNA using the following formula:

Z-score = (%EdU after siRNA − plate average %EdU) / plate standard deviation

For Skp2 and YAP nuclear/cytoplasmic ratio calculation, a four-pixel wide ring surrounding each segmented nucleus to mark the cytoplasmic region. N:C ratios for individual cells were calculated and plotted using GraphPad Prism.

#### Focal adhesion quantitative analysis

40X confocal images of phospho-FAK stained cells acquired on Operetta CLS microscope were analysed using a quantitative image-processing workflow in Fiji as described by Horzum et al^59^. Briefly, images were first pre-processed to reduce background noise and enhance contrast. Focal adhesions were then identified by intensity thresholding and segmented to generate binary masks. Individual adhesion structures were quantified using particle analysis to extract morphological parameters, including focal adhesion number, area, and size distribution.

### Cell clustering analysis

To assess the spatial clustering of the two cell populations we computed Ripley’s L-function ^60^. The L-function can be thought of as a measure of the average number of neighbouring cells within radius, *r* of a typical cell. Isotropic correction was employed to account for cells at the border of the wells as they naturally have fewer neighbouring cells. The function can be centred so that 0 represents a completely random dispersion pattern. Positive and negative values indicate clustering and repulsion between cells respectively.

To test whether the differences between Neg. C and siITGA11 conditions are statistically significant we used a permutation test originally proposed by Hahn^61^. The null hypothesis is that the point patterns across all wells are identically distributed, whereas the alternative hypothesis states that Neg. C wells share one common spatial distribution, while siITGA11 wells share another. The permutation test was performed over 3wells per condition per technical replicate containing different cell numbers. The difference in total cell numbers is accounted for since the L-function is normalised by cell density. All spatial analysis was performed using the *spatstat* https://www.routledge.com/Spatial-Point-Patterns-Methodology-and-Applications-with-R/Baddeley-Rubak-Turner/p/book/9781482210200/ package in R.

### Western blotting

Whole-cell lysates were prepared using M2 lysis buffer containing 150mM NaCl, 50 mM Tris-Cl (pH 7.4), 10% glycerol, 0.5mM EDTA, 0.5mM EGTA, and 1% Triton X-100, supplemented with phosphatase inhibitors (ThermoScientific, 1862495) and protease inhibitors (ThermoScientific, 78429). Cells were lysed on ice for 10min followed by centrifugation at 13,000 rpm for 10min.Protein concentrations were determined using the Rapid Gold BCA Protein Assay (Pierce A55860). Equal amounts of protein were mixed with 1× Novex Tris-Glycine SDS sample buffer (Novex, LC2676) supplemented with 1mM DTT and heated at 95 °C for 10min. Samples were separated on Novex 4-20% Tris-Glycine gels (Invitrogen XP04205) and transferred to PVDF-FL membranes (Merck Life Sciences IPFL00010).

Membranes were blocked in blocking buffer (TBS containing 5% milk, 10% glycerol, and 0.1% Tween-20) for 1hr at RT and incubated overnight at 4 °C with primary antibodies diluted in blocking buffer. After three washes in TBS containing 0.1% Tween-20, membranes were incubated for 1hr at RT with HRP-conjugated secondary antibodies. Membranes were washed again and developed using Clarity Western ECL substrate (Bio-Rad 1705061). Blots were imaged using an Amersham Imager 680.

Primary antibodies used included integrin α11 or ITGA11 (clone 203E1, Nanotools 0518-100, 1:1000), YAP (clone 63.7, SC101199, 1:1000), β-actin (clone 8H10D10, CST 3700, 1:1000), and vinculin (clone E1E9V, CST 13901, 1:1000). HRP-conjugated secondary antibodies were anti-mouse HRP (CST 7076P2, 1:1000) and anti-rabbit HRP (CST 7074P2, 1:1000).

### Timelapse Imaging

For all the live-cell imaging experiments, cells were plated in 384 well Phenoplates (Revvity) and imaged using an automated inverted spinning disk confocal microscope system IX83 (Olympus) set at 37 °C and 5% CO2 after every 10 min. Images were exported as tiff files and analysed using CellposeSAM segmentation^62^ and tracking using Ultrack^63^. All cells that were present throughout the entire movie were analysed unless they died. Spontaneous death was quantified manually. hTert-RPE1 mRuby-PCNA was used to define cell cycle phases as described previously^1, 35^. For timelapse imaging experiments, one experiment is shown in the results which is representative of at least two biological repeats.

### SLIC-CAGE experiment and analysis

Total RNA from two biological replicates of G0 and G1 cells was isolated using the miRNeasy Micro Kit (Qiagen) following the manufacturer’s instructions. Previously described SLIC-CAGE protocol was followed^64^, with reverse transcription, adaptor ligation, and PCR amplification for library indexing. Libraries were pooled and sequenced on Illumina NextSeq 2000.

CAGE-seq data were processed with CAGEseq v2 (https://github.com/nf-core/cageseq; cageseq2 branch, commit 07ebccd) under Nextflow v25.10.4. Reads were adapter- and quality-trimmed with Trim Galore! v0.6.7 and the unencoded 5′ guanine removed with Cutadapt v4.6, then aligned to GRCh38 (GENCODE release 45) with STAR v2.7.10a. CAGE tag start sites (CTSSs) were taken as the 5′ base of uniquely mapping read 1. In CAGEr v2.15.1 tag counts were normalised to tags per million, CTSSs below 1 TPM in every sample were discarded, and the remainder were clustered per sample (distclu, 20 bp) and aggregated across conditions into consensus clusters.

Differential expression between G0 and G1 was assessed at the level of consensus clusters. A matrix of per-replicate counts, obtained by summing the raw tag counts of the retained CTSSs within each consensus cluster, was tested with DESeq2 v1.50.2 via CAGEr’s consensusClustersDESeq2. Consensus clusters with an adjusted p-value < 0.05 and |log2 fold change| > 1 were considered differentially expressed. All analyses were performed in R v4.5.3.

### Statistical analysis

All statistical analysis used in this manuscript were performed in GraphPad Prism.

## Author contributions

ARB supervised experiments, helped to analyse data, secured funding for the work and provided feedback on the manuscript. EK, JAH and RA performed experiments and analysed data. EK prepared the figures and wrote the manuscript. AD performed cell clustering analysis, supervised by PT. ARB prepared samples for proteomics. VK, RW and TL analysed mass spectrometry data. EGM, BZ prepared samples and performed CAGE analysis, supervised by BL.

## Competing interests

The authors declare no competing interest.

## Acknowledgements

We thank colleagues in the Barr, Ly and Lenhard groups for helpful discussions. We also thank Julia Sero (University of Bath, UK) for her valuable input. We acknowledge use of the Jex HPC cluster at MRC-LMS, and the resources and support provided by the IT and Bioinformatics facilities. We thank MRC-LMS/NIHR Imperial Biomedical Research Centre Flow Cytometry Facility for support. We also thank Chad Whilding and MRC-LMS core microscopy for their help and support. We thank Wellcome Centre for Cell Biology, Edinburgh for help in mass spectrometry. ARB was supported by a CRUK Career Development Fellowship to ARB (C63833/A25729), and EK, JAH, and RA, were supported by MRC-LMS core funding (MC-A658-5TY60). VK, RW and TL were supported by Wellcome-Royal Society Sir Henry Dale Fellowship to TL (206211/A/17/Z, 218305/Z/19/Z). AD was supported by the Roth Scholarship through Department of Mathematics, Imperial College London. BZ was funded by the European Union-NextGenerationEU (grant NPOO.C3.2. R2-I1.06.0024) and the Croatian Science Foundation (project IP-2024-05-5224). PT is supported by a UKRI Future Leaders Fellowship (MR/T018429/1).

## Funder information declared

CRUK Career Development Fellowship (C63833/A25729), and MRC-LMS core funding (MC-A658-5TY60).

Wellcome-Royal Society Sir Henry Dale Fellowship (206211/A/17/Z, 218305/Z/19/Z). UKRI Future Leaders Fellowship (MR/T018429/1).

## Notes

### Competing Interest Statement

The authors have declared no competing interest.

