## Supplementary figures and legends for "Integrin α11 is enriched in quiescence and promotes cell-cycle re-entry through destabilisation of the CDK inhibitor p27"

**S1:**

A) Fluorescence-activated cell sorting (FACS) sorting of p21Low PCNALow (G1), p21High PCNALow (G0) and p21High (senescent) hTert-RPE1 mRuby-PCNA p21-GFP cell line.

B) Single-cell fluorescence intensity measurements of pRb (S807/S811) and p21-GFP of FACS sorted populations. Data shown are representative of n=3 biological repeats.

C) Representative images of β-galactosidase staining in untreated and 50μM etoposide-treated hTert-RPE1 mRuby-PCNA p21-GFP cells. Images are representative of three independent biological replicates. Scale bar =10μm.

D) Gene ontology molecular function analysis of the genes encoding the 148 proteins significantly enriched or depleted in G0/senescent cells compared to G1 cells.

E) (Left) Schematic overview of the primary image-based functional screens performed in hTert-RPE1 mRuby-PCNA p21-GFP cells under asynchronously cycling conditions. A pooled siRNA library targeting 148 candidate genes was screened. (Right) Z-scores were calculated based on the percentage of EdU-positive cells, representing actively cycling cells, for each siRNA condition. si*CCND1* was included as a positive control, producing the lowest Z-score. si*ITGA11* did not exhibit any significant effect on asynchronously cycling cells in the screen.

F) (Top) Schematic overview of the primary image-based functional screens performed in hTert-RPE1 mRuby-PCNA p21-GFP cells following two- or three-days of CDK4/6i arrest and release. A pooled siRNA library targeting 148 candidate genes was screened. (Bottom) Z-scores were calculated based on the percentage of EdU-positive cells, representing cells re-entering cell cycle, for each siRNA condition. si*CCND1* and si*TP53* were included as positive controls, producing the lowest and highest Z-scores, respectively. si*ITGA11* increased the G0 fraction (decreased EdU incorporation) following two- or three-days of CDK4/6i arrest and release.

G-I) Correlation analysis of the percentage of EdU-positive cells between two technical replicates of the siRNA screen performed in hTert-RPE1 mRuby-PCNA p21-GFP cells under G) asynchronously cycling conditions, (H) following two-days of CDK4/6i or (I) three-days of CDK4/6i arrest and release.

**S2:**

A) Percentage of EdU positive cells quantified from a 24hr EdU pulse in asynchronous, G0 (CDK4/6i), G0 (contact-inhibited) and 50μM etoposide-induced senescent hTert-RPE1 cells. Data are shown as mean ± SD (n=3 biological replicates). One-way ANOVA with multiple-comparison correction. ****P < 0.0001.

B) Western blot showing the levels of integrin $\alpha$11 in asynchronous, G0 (CDK4/6i), and C) contact-inhibited hTert-RPE1 cells transfected with either Neg.C and or si*ITGA11* for 72hrs. Vinculin is included as a loading control. Data are shown as mean ± SD (n=2 biological replicates). Unpaired student's t-test performed for statistical test, **P < 0.01.

D) Percentage of EdU positive cells positive cells quantified from a 24hr EdU pulse in asynchronous, G0 (CDK4/6i), and 50μM etoposide-induced senescent hTert-BJ fibroblast cells. Data are shown as mean ± SD (n = 3 biological replicates). One-way ANOVA with multiple-comparison correction. ****P < 0.0001.

E) (Left) Representative images and (right) quantification of integrin α11 intensity in Neg. C and si*ITGA11* transfected cells released for 24hrs from CDK4/6i-mediated arrest. Data are representative of n=3 biological replicates. Scale bar = 50μm.

F) CAGE-derived transcription from *MAX* and *EZH2* promoters in G0 and G1 cycle phases. TPM values are shown per phase; raw tag counts from the two biological replicates were pooled at the CTSS level before normalisation. Log2 fold changes (G1 versus G0) and Benjamini-Hochberg-adjusted p-values are from DESeq2 (n = 2 per phase).EZH2 marked by asterisk was found to statistically significant in differential expression analysis.

G) Schematic showing the different binding sites of *MAX* and *EZH2* factors on *ITGA11* gene as reported in the ENCODE database.

H) *MAX* and *EZH2* mRNA expression was measured by qPCR following siRNA-mediated depletion of MAX or EZH2 for 48hrs. Unpaired student's t-test performed for statistical test, ****P < 0.0001.

I) Unstained and secondary antibody controls for flow cytometric analysis of cell-surface integrin α11 expression using a Zenon™ 488-labelled anti-ITGA11 antibody. Data are representative of n=2 biological replicates.

J) Flow cytometric analysis of cell-surface integrin α11 expression using a Zenon™ 488-labelled antibody in cells induced into quiescence by 1μM CDK4/6i and transfected with either Neg. C or si*ITGA11*. Data are representative of n=2 biological replicates.

**S3:**

A) (Left) Representative images showing integrin α11 expression in different shRNA clones released from CDK4/6i-mediated arrest. Doxyxcycline (10μg/ml) was added every 24hr throughout the CDK4/6i-mediated cell-cycle arrest and continued during the release period. (Right) Quantification of cytoplasmic integrin α11 expression in different shRNA cell lines. Data is representative of the three biological replicates. Both the shRNA sequences show depletion in integrin α11 after doxycycline addition.

B) Percentage of EdU-positive in CDK4/6i-released hTert-RPE1 mRuby-PCNA shITGA11 cell lines after doxyxcycline induction (10μg/ml). Doxyxcycline was added every 24hr throughout the CDK4/6i-mediated cell-cycle arrest and continued during the release period. Quantification of EdU positive cells in different shRNA cell lines is shown. Data are shown as mean ± SD (n = 3 biological replicates with 3 technical replicates per repeat). One-way ANOVA with multiple comparison, *P < 0.05, **P < 0.01.

C) Percentage of EdU-positive cells in asynchronous hTert-RPE1 mRuby-PCNA shITGA11 cell lines following 24hrs of doxycycline induction (10µg/ml) was determined as depicted. Quantification of EdU-positive cells across different shRNA cell lines is shown. Data are shown as mean ± SD (n = 3 biological replicates with 3 technical replicates per repeat). One-way ANOVA with multiple comparison, none of the comparisons were found to be statistically significant.

**S4:**

A) (Left) Representative images of hTert-RPE1 mRuby-PCNA cells transfected with Neg. C or si*ITGA11*, showing cell clustering 24hrs post CDK4/6i-release. Red dots indicate image acquisition during CDK4/6i-mediated arrest and green dot indicates CDK4/6i-released time point following Hoechst 33582 staining. (Right) Graph depicts the coefficient of variation of Ripley's L-function as a function of radius and is generated from n=3 biological replicates. Permutation test originally proposed by Hahn[59] was used for statistical analysis.

B) Percentage of EdU-positive hTert-RPE1 mRuby-PCNA cells transfected with Neg. C or si*ITGA11* in asynchronous cells. Contact-inhibited hTert-RPE1 mRuby-PCNA cells were included as a control to confirm efficient induction of quiescence, as demonstrated by the marked reduction in EdU-positive cells. Data are presented as mean ± SD (n = 3 biological replicates). One-way ANOVA with multiple comparison, differences between Neg.C and si*ITGA11* in asynchronous cells were not significant. ****P < 0.0001.

C) (Left) Representative images depicting cells after contact inhibition and re-seeding of hTert-RPE1 mRuby-PCNA cells transfected with Neg. C and si*ITGA11*. Red dots indicate image acquisition after replating and reverse transfection of contact inhibited cells in serum free media and green dot indicates cells released into complete media following Hoechst 33582 staining. (Right) Graph shows the coefficient of variation in Ripley's L-function across different radii at 24hrs post release of contact inhibited cells in complete media (green dot). Permutation test originally proposed by Hahn[59] was used for statistical analysis. Plot represents data from n=3 biological replicates.

D) (Left) Graph shows total area of focal adhesions, as determined from pFAK immunostaining in hTert-RPE1 mRuby-PCNA cells transfected with Neg. C or si*ITGA11* following 24hrs release from CDK4/6i. (Right) Images representing phospho-FAK and Hoechst 33582 staining from Neg. C and si*ITGA11* are shown. Representative focal adhesions are colour coded in red. Data shown as mean ± SD is representative of n=2 biological repeats. Scale bar = 20µm.

**S5**:

A) Graphs show cell cycle phases in asynchronously growing hTert-RPE1 mRuby-PCNA cells as monitored by live-cell imaging 6hrs after siRNA transfection. Each row represents an individual cell.

B) Quantification of G1, S, G2 and M phase lengths in Neg. C and si*ITGA11* conditions is shown. Data is representative of the two biological repeats. Unpaired student's t-test performed for comparison and was not significant.

C) Percentage of cells undergoing cell death in hTert-RPE1 mRuby-PCNA cells 6hrs after transfection with either Neg. C or si*ITGA11*. Unpaired student's t-test performed for comparison, *P < 0.05.

**S6:**

A) Representative images of p27 immunostaining in hTert-RPE1 mRuby-PCNA CDK4/6i released cells transfected with Neg.C or si*ITGA11* for 72hrs before fixation and staining. Scale bar=100μm.

B) (Left) Quantification of nuclear p27 intensity in contact-inhibited hTert-RPE1 mRuby-PCNA cells. Cells were released from contact inhibition by re-seeding and transfected with Neg.C or si*ITGA11* at the time of re-seeding (24hrs after cell cycle re-entry). Nuclear p27 intensity was quantified 48hrs after re-seeding/transfection. Data are presented as mean ± SD (n = 2 biological replicates). Unpaired student's t-test performed for comparison, ****P < 0.0001. (Right) Representative images of p27 immunostaining in hTert-RPE1 mRuby-PCNA CDK4/6i released cells released from contact inhibition by re-seeding and transfected with Neg.C or si*ITGA11* for 72hrs before fixation and staining. Scale bar=100μm.

C) Quantification of nuclear p27 fluorescence intensity in CDK4/6i-released hTert-RPE1 mRuby-PCNA shRNA cell lines 24hrs after doxycycline induction (10μg/ml). Data shown as mean ± SD (n = 3 biological repeats). One-way ANOVA statistical test performed for multiple comparisons, ****P < 0.0001.

D) Quantification of nuclear p27 fluorescence intensity in asynchronous hTert-RPE1 mRuby-PCNA shRNA cell lines 24hrs after doxycycline induction (10μg/ml). Data shown as mean ± SD (n = 3 biological repeats). One-way ANOVA statistical test performed for multiple comparisons, ****P < 0.0001.

E) Quantification of nuclear Cyclin D1 fluorescence intensity in CDK4/6i-released hTert-RPE1 mRuby-PCNA cells transfected with Neg. C or si*ITGA11*. Data are presented as mean ± SD (n = 2 biological replicates). Unpaired student's t-test performed for comparison, ****P < 0.0001.

F) Quantification of nuclear p27 fluorescence intensity in (left) asynchronous and (right) CDK4/6i-released hTert-RPE1 mRuby-PCNA cells transfected with Neg. C or si*CDKN1B* to show efficiency of siRNA. Data are presented as mean ± SD (n = 3 biological replicates). Unpaired student's t-test performed for comparison, ****P < 0.0001.

G) Representative images of Skp2 immunostaining in hTert-RPE1 mRuby-PCNA CDK4/6i released cells transfected with Neg.C or si*ITGA11* and released for 24hrs before fixation and staining. Scale bar=100μm.

**S7:**

A) *CTGF*, *CYR61* and *ITGA11* mRNA expression in hTert-RPE1 mRuby-PCNA CDK4/6i released cells transfected with Neg.C or si*ITGA11* for 72hrs. *GAPDH* was used as a reference gene. Data shown as mean ± SD (n = 4 biological repeats). Unpaired T-test performed for comparison, **P < 0.01.

B) *CTGF*, *CYR61* and *ITGA11* mRNA expression in hTert-RPE1 mRuby-PCNA CDK4/6i released cells transfected with Neg.C or si*ITGA11* for 72hrs with 3μM LATS1/2 inhibitor (TDI-011536) or DMSO. GAPDH was used as a reference gene. Data shown as mean ± SD (n = 3 biological repeats). One-way ANOVA statistical test performed for multiple comparisons, *P<0.05, **P < 0.01, ***P<0.001, ****P < 0.0001.

C) (Left) Western blot showing the expression of phospho-YAP (Ser127), total YAP and integrin α11 in hTert-RPE1 mRuby-PCNA CDK4/6i released cells transfected with Neg.C or si*ITGA11* for 72hrs with 3μM LATS1/2 inhibitor (TDI-011536) or DMSO. (Right) Quantification of the phospho-YAP and total YAP normalized to β-actin. Data are presented as mean ± SD from two biological replicates. The specific band of YAP is marked as asterisk. Statistical significance was determined by one-way ANOVA with multiple-comparison correction. ****P < 0.0001.

D) Quantification of nuclear p27 fluorescence intensity in CDK4/6i-released hTert-RPE1 mRuby-PCNA cells transfected with Neg. C or si*ITGA11* for 72hrs with 3μM LATS1/2 inhibitor (TDI-011536) or DMSO. Data are shown as mean ± SD and is representative of n = 2 biological repeats. Statistical significance was determined by one-way ANOVA with multiple-comparison correction. ****P < 0.0001.

E) Quantification of nuclear Skp2 fluorescence intensity in CDK4/6i-released hTert-RPE1 mRuby-PCNA cells transfected with Neg. C or si*ITGA11* for 72hrs with 3μM LATS1/2 inhibitor (TDI-011536) or DMSO. Data are shown as mean ± SD and is representative of n = 2 biological repeats. Statistical significance was determined by one-way ANOVA with multiple-comparison correction. ****P < 0.0001.

**Supplementary tables**

Supplementary table 1: List of proteins identified from LC-MS. Normalized raw intensities for each of these proteins for p21Low PCNALow G1 cells (Lo), p21High PCNALow spontaneously G0 (Hi) and p21High senescent cells (Sen) are shown for three biological replicates.

Supplementary table 2: List of the 148 candidate proteins selected for the image-based siRNA screen, showing their representative clusters identified by quantitative LC-MS analysis.

Supplementary table 3: Percentage of EdU positive cells in hTert- RPE1 cells released from two-day CDK4/6 inhibitor treatment and transfected with ONTarget siRNA Pools (ThermoFisher) targeting 129 genes including controls.

Supplementary table 4: Percentage of EdU positive cells in hTert-RPE1 cells released from three-day CDK4/6 inhibitor treatment and transfected with ONTarget siRNA Pools (ThermoFisher) targeting 129 genes including controls.

Supplementary table 5: Percentage of EdU positive cells in asynchronously cycling hTert-RPE1 cells transfected with ONTarget siRNA Pools (ThermoFisher) targeting 129 genes including controls for 48hrs.

Supplementary table 6: List of genes whose depletion decreased EdU positive or G1 fractions in asynchronously cycling cells.

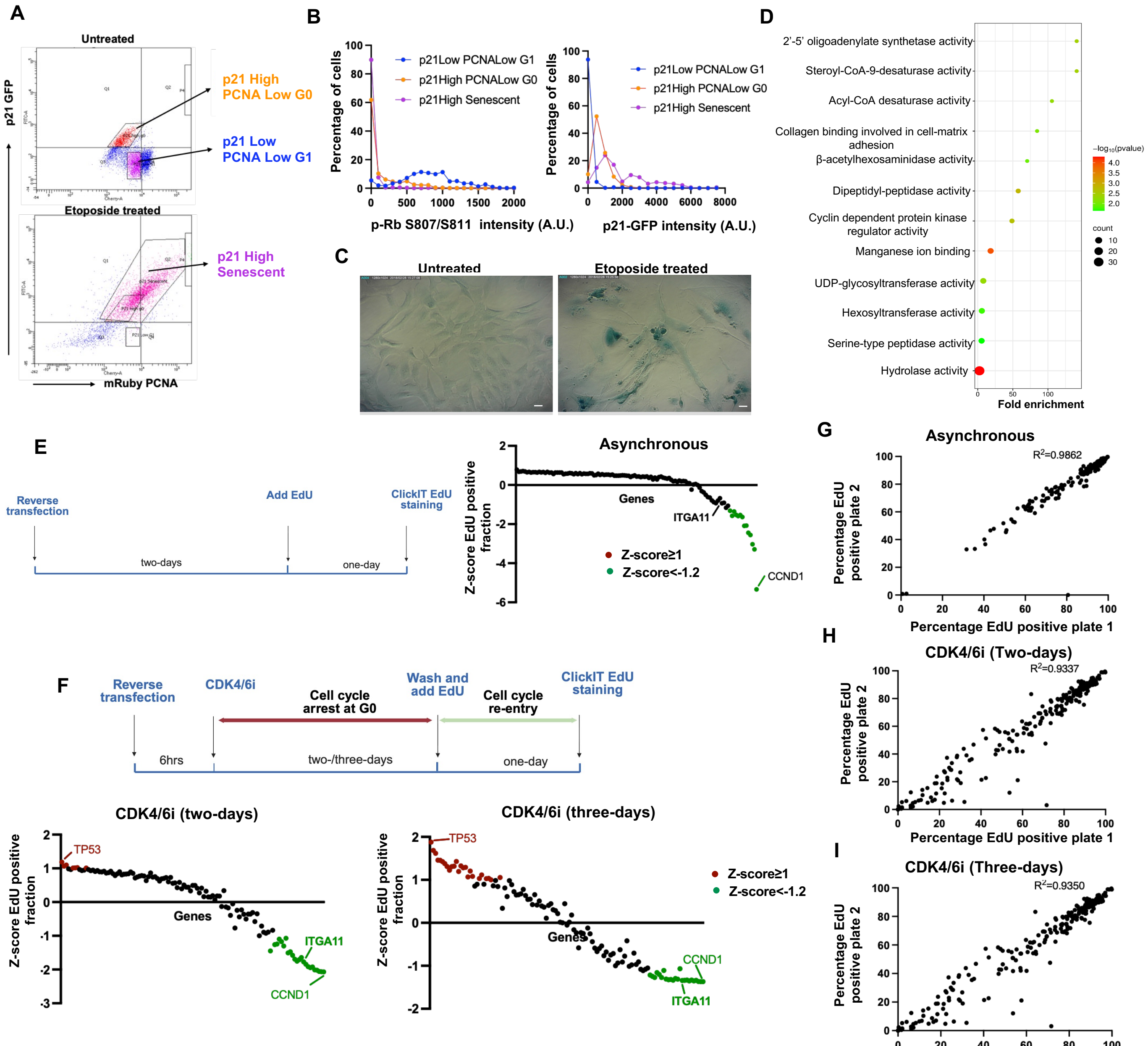


Supplementary Figure 1

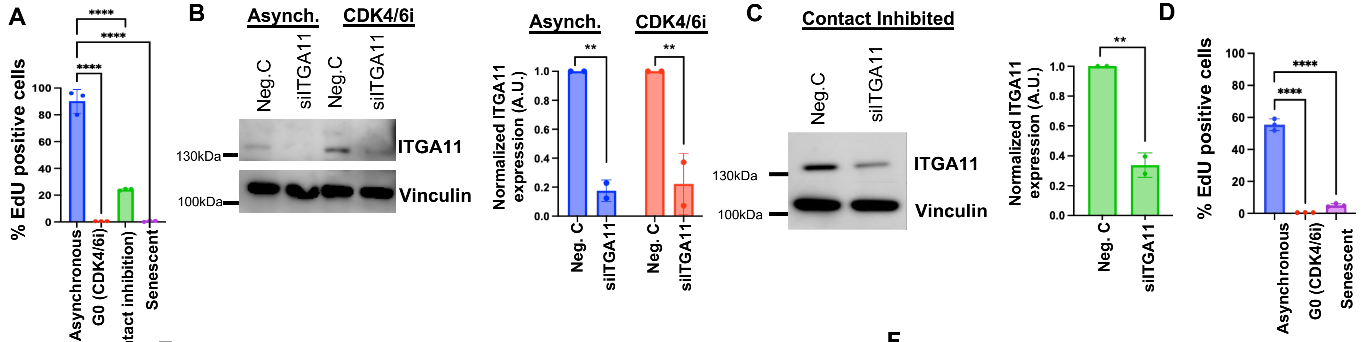


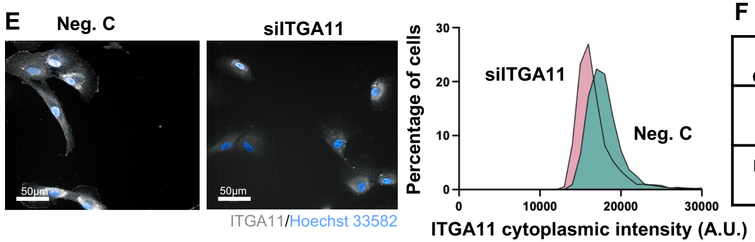


F

| Genes | Chromosomal locations | G0 | G1 | Log2 Fold Change | padj |
|---|---|---|---|---|---|
| MAX | Chr14: 65102486-65102539 | 33.29 | 23.05 | -0.44 | 0.235 |
| EZH2* | Ch7: 148884287-48884292 | 1.44 | 6.48 | 2.25 | 2.66E-09 |

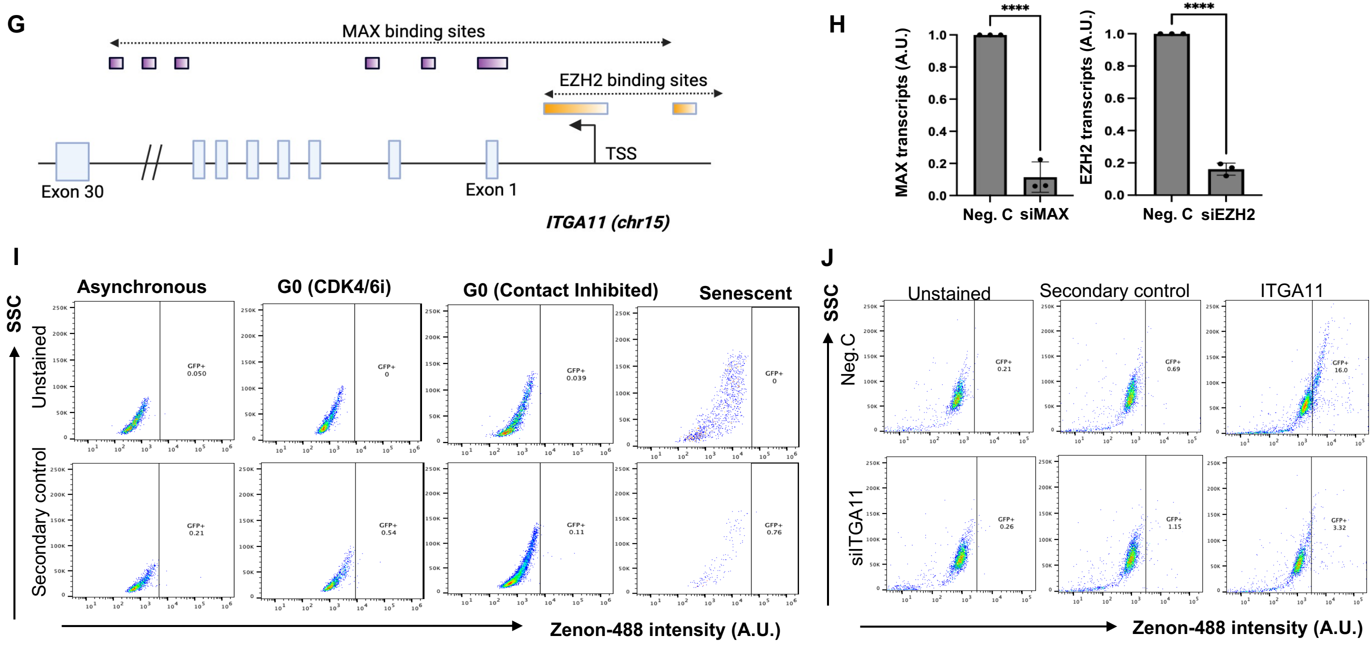


Supplementary Figure 2

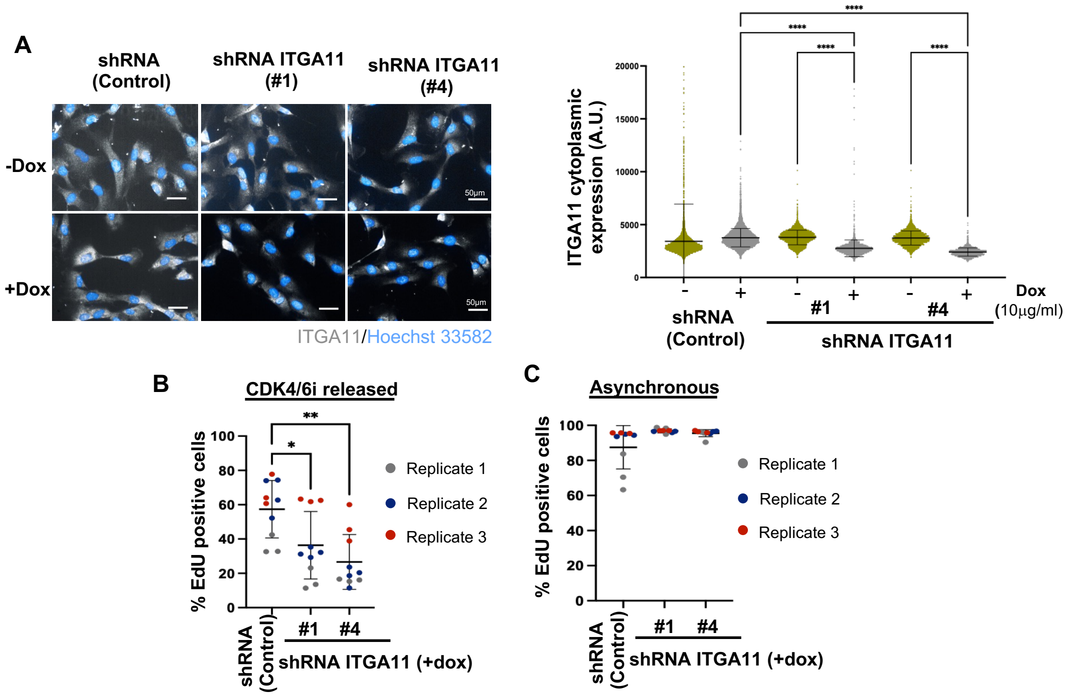


Supplementary Figure 3

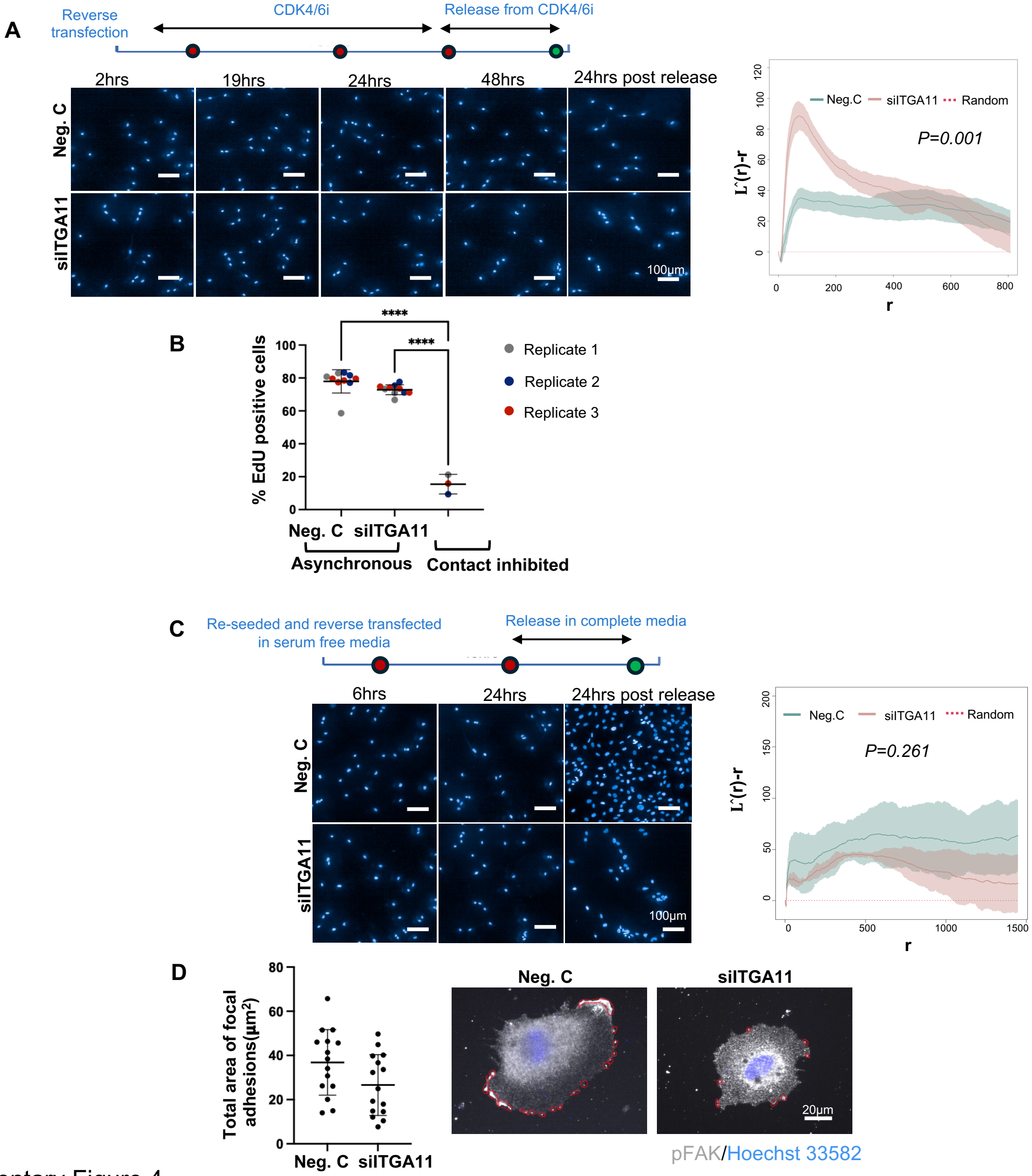


Supplementary Figure 4

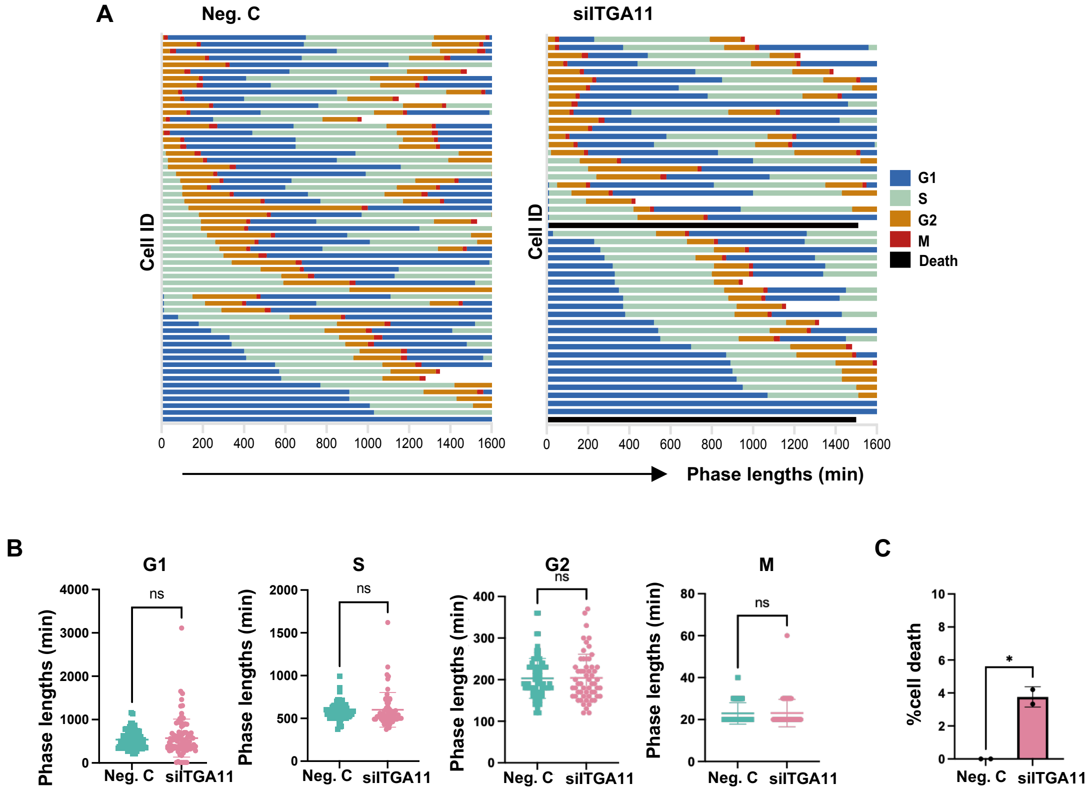


Supplementary Figure 5

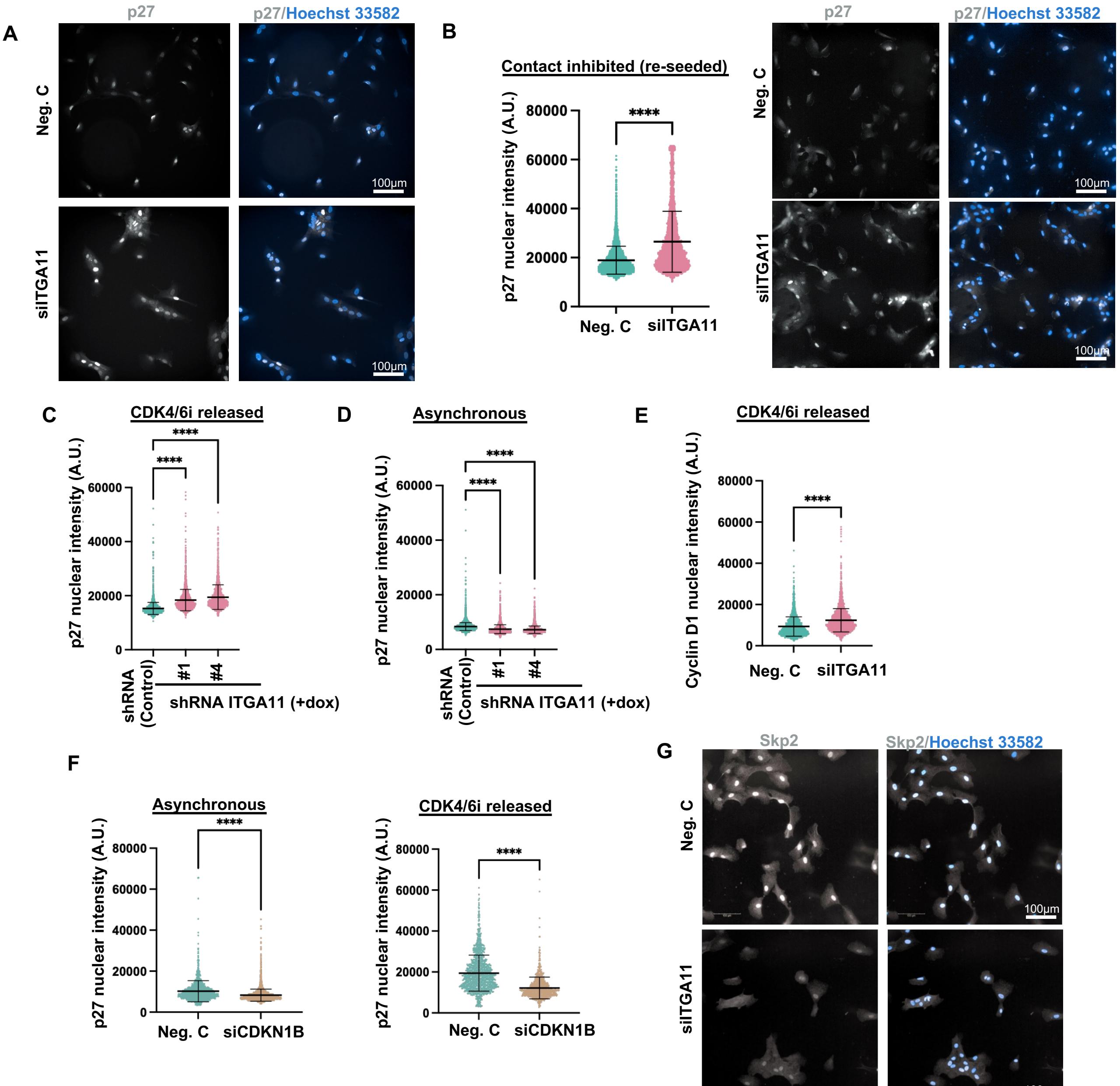


Supplementary Figure 6

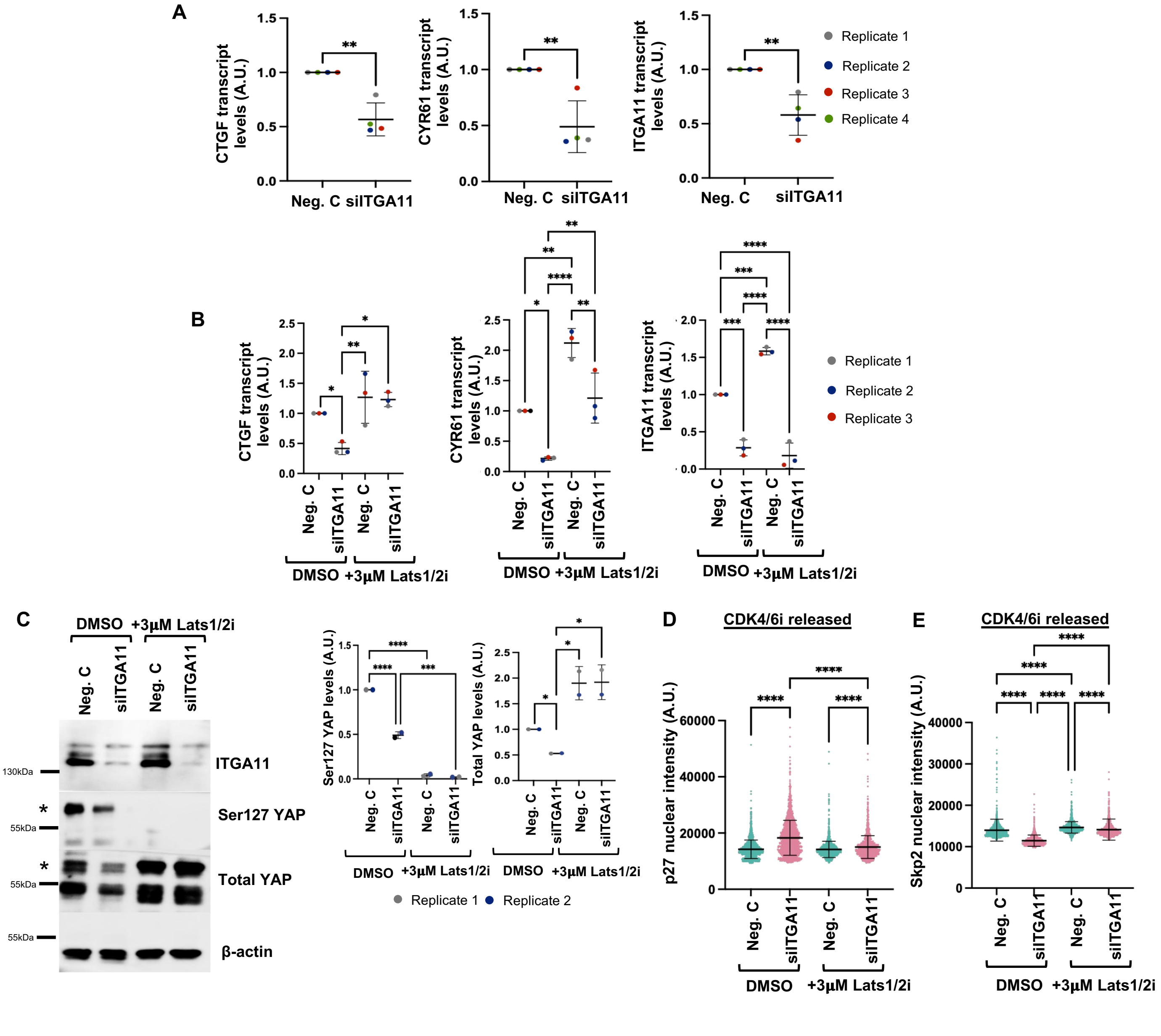


Supplementary Figure 7
